# Replication-competent SIVcpz CRISPR screen identifies barriers to successful cross-species transmission

**DOI:** 10.1101/2025.10.27.684817

**Authors:** Qinya Xie, Qingxing Wang, Sabrina Noettger, Guillermo Gosálbez, Annika Betzler, Meta Volcic, Dorota Kmiec, Stefan Krebs, Alexander Graf, Dila Gülensoy, Gilbert Weidinger, Konstantin M.J. Sparrer, Frank Kirchhoff

**Author notes:** These authors contributed equally to this work (Qinya Xie, Qingxing Wang, Sabrina Noettger, Guillermo Gosálbez).

## Abstract

Simian immunodeficiency viruses (SIVs) have crossed from apes to humans at least four times, but only one event gave rise to the AIDS pandemic. The host barriers that pandemic HIV-1 group M (*major*) strains overcame to spread efficiently in humans remain poorly understood. To identify such barriers, we performed CRISPR-Cas9 screens driven by the replication efficiency of SIVcpz, the chimpanzee precursor of HIV-1. Guide RNA libraries targeting more than 500 human genes encoding potential antiviral factors were inserted into the replication-competent SIVcpz MB897 molecular clone, which is phylogenetically closely related to HIV-1 group M strains. Propagation in Cas9-expressing human SupT1 T cells significantly enriched for sgRNAs targeting ADAR, *AXIN1, CEACAM3, CD72, EHMT2, GRN, HMOX1, HMGA1, ICAM2, CD72, IFITM2, MEFV, PCED1B, SGOL2, SMARCA4, SUMO1* and *TMEM173*. These hits only partially overlapped with those identified in analogous HIV-1–based screens, indicating virus-specific restriction profiles. Functional analyses confirmed that IFITM2 (interferon-induced transmembrane protein 2), PCED1B (PC-esterase domain–containing protein 1B), MEFV (Mediterranean fever protein, pyrin/TRIM20), and AXIN1 (Axis inhibition protein 1), restrict replication of SIVcpz but not of HIV-1 group M strains in primary human CD4⁺ T cells. These findings reveal previously unrecognized host factors that limit SIVcpz replication in human cells and highlight barriers that HIV-1 likely overcame during its adaptation for pandemic spread.

**One Sentence Summary:** CRISPR screens with replication-competent SIVcpz identify human antiviral factors limiting efficient viral replication after zoonotic transmission.

## INTRODUCTION

Viral pathogens and their hosts are engaged in a continuous evolutionary arms race (*1*). Functionally and structurally diverse host-encoded antiviral factors provide rapid, broad-spectrum protection against infection (*2–4*). In response, many viruses evolved strategies to evade or counteract these defenses, leading to a dynamic interplay that shapes infection outcomes (*3*, *5*, *6*). Failures in antiviral host defense can result in severe disease and - in very rare cases - global pandemics (*7*, *8*).

HIV-1, the causative agent of AIDS, emerged through four cross-species transmissions of simian immunodeficiency viruses infecting chimpanzees (SIVcpz) and western lowland gorillas (SIVgor) (*8*, *9*). However, only one transmission of SIVcpz to humans, originating from a central chimpanzee (*Pan troglodytes troglodytes, Ptt*) about 100 years ago that gave rise to the M (*major*) group of HIV-1 is responsible for the AIDS pandemic (*9*). In contrast, SIVcpz strains from eastern chimpanzees (*Pan troglodytes schweinfurthii, Pts*) have not been detected in humans (*10*), indicating the importance of both viral and host-specific factors for successful zoonotic transmission. One reason for the efficient spread of HIV-1 group M strains is their ability to counteract human restriction factors. For example, their accessory protein Vpu evolved to antagonize tetherin (BST-2), a host factor that inhibits virion release (*11*, *12*). Because HIV-1 group M strains are already well adapted to the human host and effectively evade innate immune defenses, they are poorly suited for identifying the antiviral mechanisms that were overcome during their emergence.

To identify antiviral defenses that posed early barriers to efficient spread of HIV-1 in humans, we adapted our previously developed CRISPR-based screening approach (*13*) by replacing pandemic HIV-1 group M constructs with a closely related SIVcpz*Ptt* strain. Unlike HIV-1 group M strains, SIVcpz is not adapted to human cells and thus provides a useful tool for identifying antiviral mechanisms that were overcome during HIV-1 adaptation. These factors may continue to limit lentiviral zoonoses and restrict replication of less-adapted HIV-1 lineages, including group O, N, and P viruses. We employed the “traitor virus” (TV) approach, in which replication-competent infectious molecular clones (IMCs) are engineered to express single guide RNAs (sgRNAs) targeting host genes. In Cas9-expressing T cells, viruses encoding sgRNAs against antiviral genes gain a replication advantage, enabling identification of restriction factors based on sgRNA enrichment during serial viral passaging (*13*). Unlike conventional overexpression, RNAi, or single-cycle CRISPR screens, the TV strategy does not require manipulation of cellular factors prior to infection cells and is driven by viral replication efficacy. This allows discovery of host genes with modest effects in single-round assays but substantial impact across multiple replication cycles, better covering the entire viral replication cycle, and reflecting the dynamics of viral spread *in vivo*.

We previously used libraries of replication-competent HIV-1 constructs encoding >1500 sgRNAs for virus-driven discovery of antiviral factors (*13*). Here, we extended this technique to SIVcpz*Ptt* to uncover antiviral factors and mechanism that pandemic HIV-1 strains evolved to counteract or evade during adaptation to the human host. Propagation of SIVcpz*Ptt* in Cas9 expressing SupT1-R5 cells led to the enrichment of sgRNAs targeting host factors that overlapped with but were distinct from those previously identified in HIV-1-based screens. Functional analyses confirmed that several of these factors restrict SIVcpz*Ptt* but not HIV-1 in primary human CD4+ T cells. Our results reveal barriers to successful lentiviral zoonoses and provide insights into the host determinants that shaped the emergence and pandemic potential of HIV-1.

## RESULTS

### SIVcpz*Ptt* MB897 replicates efficiently in SupT1-R5 Cas9 cells

To uncover host factors that restricted early spread of HIV-1, we performed a CRISPR-based screen using the SIVcpz*Ptt* MB897 infectious molecular clone (IMC). This strain was selected because it is closely related to pandemic HIV-1 M strains, including the NL4-3 and CH077 IMCs used in our initial screen (Fig. 1A), and replicates in human cells and humanized mice (*14, 15*, *16*).

**Fig. 1.**
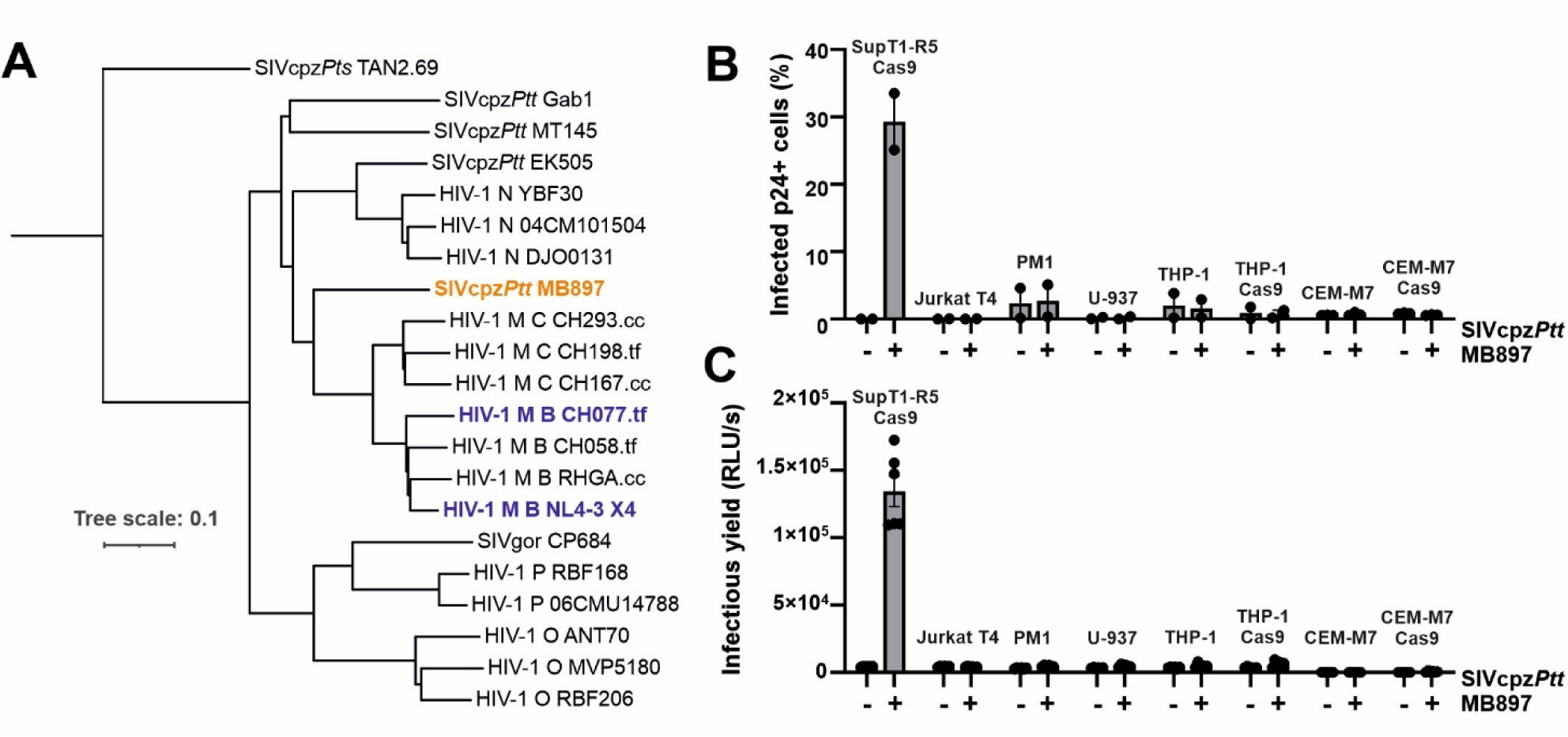
Phylogenetic relationship and infectivity of SIVcpz*Ptt* MB897. (**A)** Evolutionary relationship of SIVcpz and SIVgor strains with M, N, O, and P strains of HIV-1. The tree was constructed based on the genomic sequences using the maximum likelihood method (*21*). Scale bar: 0.1 nucleotide replacements per site. The SIVcpz*Ptt* MB897, HIV-1 CH077 and NL4-3 IMCs used in the present and previous screens are highlighted in orange and blue, respectively. Tf, transmitted- founder; cc, chronic control. (**B)** Percentages of cells productively infected with SIVcpz*Ptt* MB897 (p24+ and CD4 low). Infection was determined by flow cytometric analysis at three days post- infection (Fig. S1). (**C**) Infectious SIVcpz*Ptt* MB897 production was determined by infecting TZM- bl reporter cells with supernatants obtained from the indicated cell lines 5 days after virus exposure.

To identify suitable human cell lines for virus propagation, we generated SIVcpz*Ptt* MB897 stocks by transfection of 293T cells and used them to infect SupT1-R5, Jurkat, PM1, U-937, THP-1 and CEM-M7 cells. SupT1-R5 (modified to express CCR5) and Jurkat cells were derived from T-cell leukemias (*16*, *17*), and PM1 cells from peripheral blood lymphocytes (*18*). CEM-M7 is a CEM subclone with enhanced susceptibility to interferon (IFN) (*19*). U-937 and THP-1 are monocytic leukemia lines that can differentiate into macrophage-like cells. We previously generated Cas9-expressing derivatives of SupT1-R5 and CEM-M7 cells (*13*).

Productive infection was assessed by flow-cytometric analysis of intracellular p24 capsid antigen expression and CD4 down-modulation by Nef (Fig. S1). We found that only SupT1-R5 Cas9 cells were susceptible to SIVcpz*Ptt* MB897 infection (Fig. 1B). To further validate this, supernatants from all cell cultures were used to infect TZM-bl cells, which express CD4, CCR5, as well as CXCR4, and contain Tat-responsive β-galactosidase and luciferase reporters (*20*). Supernatants from SupT1-R5 Cas9 cells induced high β-galactosidase activity (Fig. 1C), indicating effective production of infectious SIVcpz*Ptt* MB897. We therefore used SupT1-R5 Cas9 cells in subsequent experiments.

### Generation of sgRNA-expressing SIVcpz*Ptt* MB897 constructs

To identify antiviral cellular genes, we engineered replication-competent SIVcpz*Ptt* MB897 constructs carrying a compact (∼351 nt) sgRNA expression cassette. As previously reported (*13*), this cassette contains the human U6 promoter and an sgRNA with a flexible targeting sequence and an invariant scaffold. All viral genes remain intact and are expressed under the native LTR and splice sites, while sgRNAs are independently transcribed from the U6 promoter (Fig. 2A). To preserve a functional *nef* gene, we eliminated its overlap with the U3 region of the viral 3’LTR and inserted the sgRNA-cassette between them. Previous analyses of HIV-1 constructs showed that duplicated *nef*/U3 sequences or direct repeats may drive loss of the cassette during viral replication (*13*). To circumvent this, we designed three SIVcpz*Ptt* variants: (1) Mut1, with codon-optimized *nef* gene to minimize homology with the U3 region; (2) Mut2, that harbors a truncated 3’LTR U3 region but maintains critical *cis-acting elements* and the core enhancer/promoter elements. Notably, large parts of the U3 region serve mainly as Nef coding region and are dispensable for viral replication both *in vitro* and *in vivo* (*22–24*). (3) Mut3, is similar to Mut2 but contains additional changes at the 3’end of *nef* to remove another potential recombination site (Fig. 2A).

**Fig. 2.**
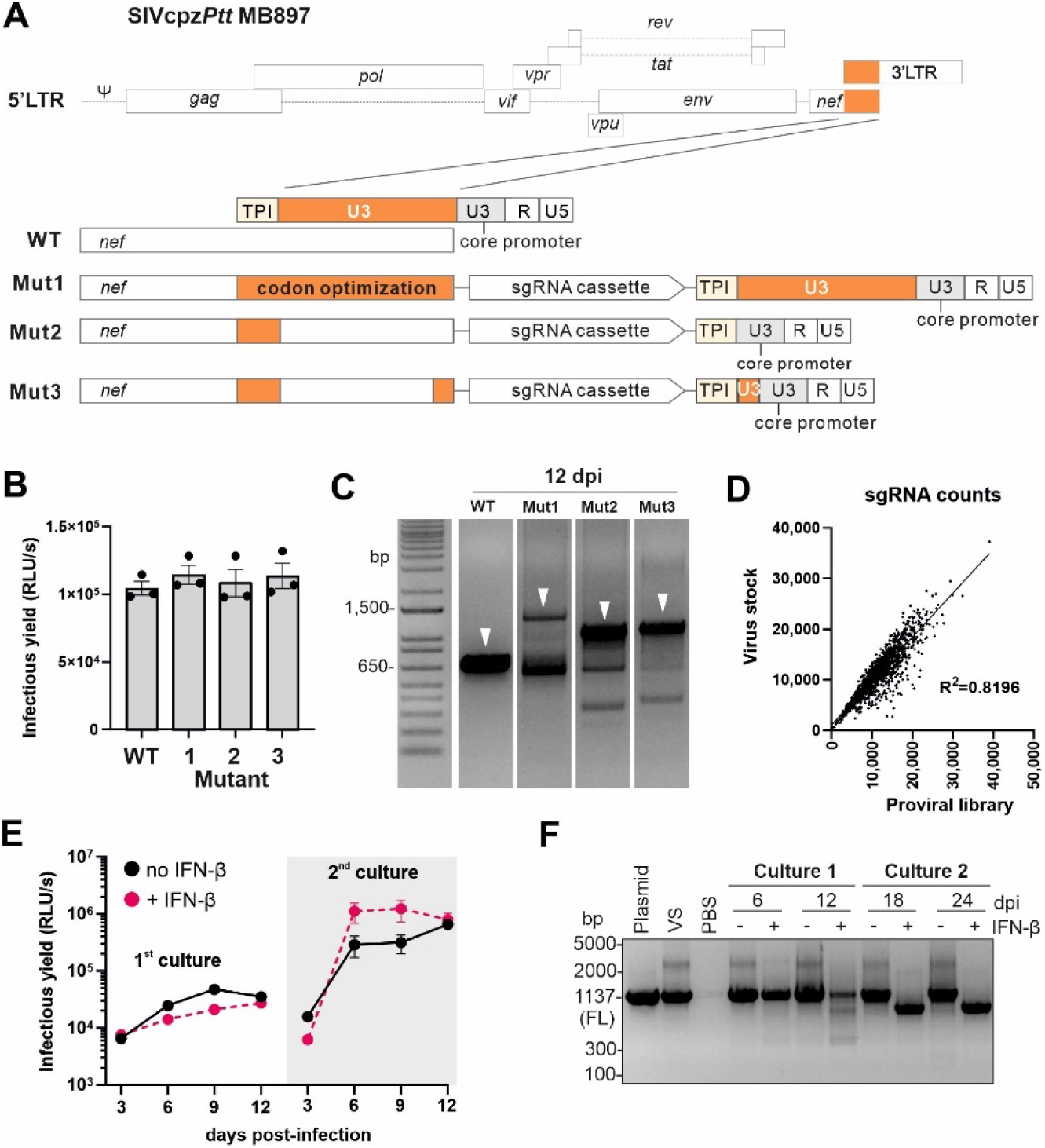
Generation and characterization of SIVcpz*Ptt* MB897 sgRNA constructs. (**A**) Schematic of sgRNA expressing SIVcpz*Ptt* MB897 IMCs. The sgRNA expression cassette was inserted between *nef* and the 3’LTR. Mut1 contains a duplicated nef/3’LTR overlap with codon-optimized *nef*. Mut2 and Mut3 lack parts of U3 that serve mainly as Nef coding region but maintain critical cis regulatory element: a T-rich region, the polypurine tract, and attachment sites required for integration, named the TPI region. (**B**) Infectious virus yield from HEK293T cells transfected with parental or Mut1–Mut3 constructs, measured by TZM-bl reporter assay (mean ± SD of three experiments). (**C**) Stability of the constructs in SupT1-CCR5- Cas9 cells, assessed by RT-PCR 12 dpi (days post-infection). Arrows indicate expected fragment sizes. (**D**) Correlation between sgRNA read counts in viral stocks and proviral DNA from the MB897 sgRNA library. (**E**) Infectious virus production in SupT1-R5 Cas9 cells infected with the SIVcpz*Ptt* MB897mut3-sgRNA library, passaged as indicated in Fig. S2. Virus titers in supernatants were measured by TZM-bl assay. (**F**) RT-PCR analysis of MB897mut3-sgRNA constructs in the corresponding supernatants. FL, full-length.

Transfection of all proviral constructs yielded high levels of infectious SIVcpz*Ptt* MB897 (Fig. 2B), which replicated efficiently in SupT1 R5 Cas9 cells. However, RT-PCR analyses revealed that Mut1 largely lost the U6-sgRNA-scaffold cassette after 12 days, whereas Mut2 and Mut3 remained stable (Fig. 2C). Since Mut3 showed only minimal cassette loss, we selected it for generation of the SIVcpz-based library. As reported for HIV-1 (13), we generated SIVcpzPtt TV constructs targeting 510 candidate antiviral genes, each by three distinct sgRNAs. Of these, 200 genes shared features of known antiviral factors (25), and the remaining 310 were chosen because of their potential roles in viral sensing or HIV-1 replication (26). Cloning was highly efficient and deep sequencing confirmed the presence of all 1,537 sgRNAs. Moreover, sgRNA frequencies in viral stocks closely matched those in the proviral TV-MB897-sgRNA library (Fig. 2D).

Since many well-known antiviral factors are IFN-inducible, we treated SupT1-R5 Cas9 cells with IFN-α or IFN-β. Both robustly induced ISG15, confirming activation of IFN-stimulated genes (ISGs, Fig. S3A). To identify both constitutive and IFN-induced antiviral factors, we initially cultured the TV-MB897-sgRNA library in the presence or absence of nontoxic concentrations of IFN-β (100 U/ml; Fig. S3B) for 12 days (each in two independent experiments). We then used the supernatants obtained at the end to initiate a second virus propagation period (Fig. S2). Conditions were modified compared to previous passaging of HIV-1 (*13*), because SIVcpz replicates with slower kinetics and has been reported to be more dependent on cell-to-cell spread (*27*, *28*). Infection of TZM-bl cells with culture supernatants revealed higher infectious virus production during the 2^nd^ culture period (Fig. 2E), suggesting the potential emergence of SIVcpz variants with increased replication fitness. Unexpectedly, however, SIVcpz*Ptt* TV-MB897-sgRNA constructs consistently lost the sgRNA cassette in the presence of IFN-β at later time points (Fig. 2F). In contrast, they replicated efficiently and remained stable in the absence of IFN-β. Consequently, subsequent selection experiments and deep sequencing analyses were performed in the absence of IFN.

### Enrichment of sgRNAs targeting genes restricting SIVcpz*Ptt* replication

To identify sgRNAs that promote SIVcpz*Ptt* replication, we analyzed virus containing cell culture supernatants obtained in 6-day intervals from two independent cultures by next generation sequencing (NGS). MAGeCK analysis (*29*) allowed us to follow the frequency of all sgRNAs during virus propagation. Widening volcano plots revealed progressive enrichment or depletion of specific sgRNAs (Fig. 3A) with highly significant correlations across different timepoints (Fig. 3B). Although some top scoring hits varied between early and late time points several candidates consistently emerged. At days 6 and 12, the most enriched genes included *IFITM2, PCED1B, AXIN1, CEACAM3,* and *MEFV*. At days 18 and 24, the top hits included SGOL2 (chromosome segregation) (*30*), SMARCA4 (chromatin remodeling) (*31*), TMEM173 (STING, a cytosolic DNA sensor) (*32*), and HMOX1 (a heme-degrading enzyme with anti-inflammatory effects). In agreement with the previous HIV-1 screen (*13*), sgRNAs varied in selection efficiency, but their relative effects on viral replication fitness were highly reproducible and in most cases at least two of three sgRNAs clearly enriched (Fig. 3C, S4A). Altogether, hit factors may restrict SIVcpz through diverse mechanisms including modulation of innate immunity signaling and oxidative stress, interference with viral transcription and regulation of cell cycling.

**Fig. 3.**
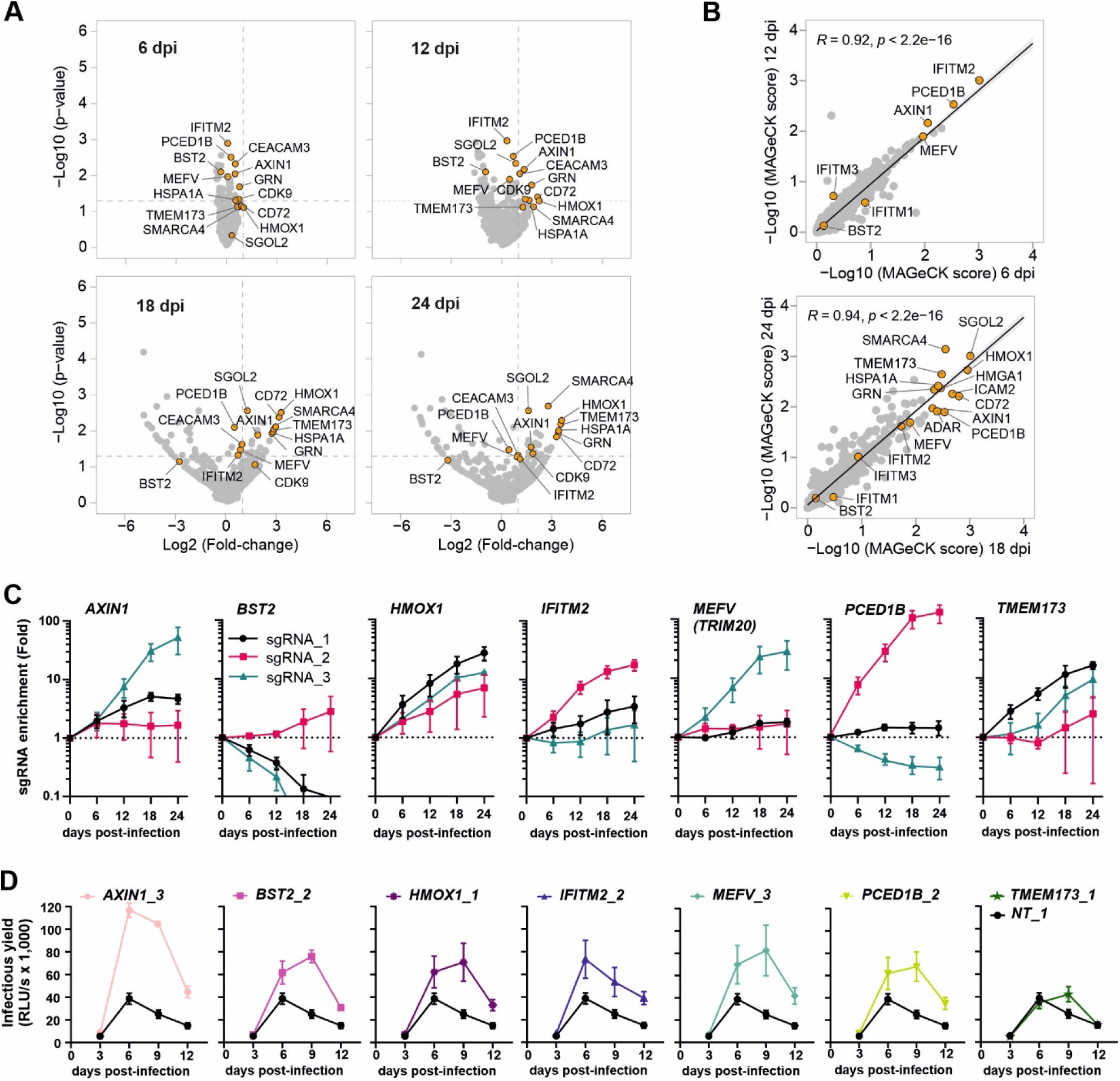
SIVcpz-driven enrichment and validation of sgRNAs. (**A**) Volcano plots indicating enrichment of specific genes targeted by sgRNAs during culture of the MB897-sgRNA library in SupT1-R5 Cas9 cells. (**B**) Correlation of gene-level MAGeCK enrichment scores at the indicated time-points post-infection. (**C**) Relative read counts from MAGeCK analysis showing enrichment of sgRNAs targeting *AXIN1, BST-2, HMOX1, IFITM2, MEFV, PCED1B* or *TMEM173* during library propagation. (**D**) Validation of individual SIVcpz*Ptt* MB897 constructs expressing targeting sgRNAs or a non-targeting control in SupT1-CCR5 Cas9 cells. Numbers after the underscore specify the sgRNA. Infectious virus yield was quantified by TZM-bl assay. Data represent means of three independent experiments ± SEM.

To validate candidate antiviral factors, we generated 8 individual SIVcpz*Ptt* MB897 constructs encoding sgRNAs targeting *AXIN1, BST-2, HMOX1, IFITM2, MEFV, PCED1B or TMEM173,* or a non-targeting (NT) control. Except for BST-2 (tetherin) these targets were chosen based on their strong enrichment in the screens (Fig. 3A-C). BST-2 (tetherin), which SIVcpz cannot antagonize in human cells (*11*, *12*), was not a significant hit because only one of three sgRNA was enriched (Fig. 3C). All targeting sgRNAs increased SIVcpz*Ptt* MB897 replication in SupT1-R5 Cas9 cells, albeit with different efficiency (Fig. 3D). While most sgRNAs enhanced infectious virus yields by ∼2-fold, AXIN1 had very strong and TMEM173 (encoding STING) only subtle and enhancing effects. Importantly, none of the sgRNAs significantly affected replication in parental SupT1-R5 cells lacking Cas9 (Fig. S4B). Altogether, these results show that sgRNAs targeting *AXIN1, BST-2, HMOX1, IFITM2, MEFV* and *PCED1B* clearly enhance SIVcpz*Ptt* replication in a Cas9/CRISPR-dependent manner in the human SupT1 T cell line.

### Overlapping sets of sgRNAs increase replication fitness of SIVcpz and HIV-1

To compare the SIVcpz-driven screen with previous HIV-1 NL4-3 and CH077 datasets using the same sgRNAs library in CEM-M7 Cas9 cells (*13*), we performed correlation analyses. As expected, the strongest correlations arose between matching viral libraries at similar time points (Fig. S5). Several sgRNAs, including *AXIN1, SUMO1, FOXP3, PRAMEF16, CD72, NDUFS5, HSPA1A, EHMT2, NFATC1, TAGLN2, GBP4 and RAD18,* displayed consistent MAGeCK scores across SIVcpz and HIV-1, while others showed virus-specific enrichment (Fig. 4A). AXIN1 is a negative regulator of Wnt signaling (*33*). SUMO1 modulates numerous cellular processes and is known to enhance TRIM5α-mediated retroviral restriction (*34*). FOXP3, a master transcriptional regulator of regulatory T cells, has been reported to exert both inhibitory and enhancing effects on HIV-1 LTR– driven gene expression (*35*, *36*). HSPA1A (Hsp70) had been reported to prevent Vpr-induced cell cycle arrest (*37*), suppress LTR transcription by displacing NF-κB p50/p65 (*38*), and to inhibit Vif-mediated ubiquitination and degradation of APOBEC3G (*39*). Notably *HSPA1B*, a close relative of *HSPA1A* also emerged in the HIV-1-driven screen (Fig. 4A, right) EHMT2 (G9a) mediates H3K9 dimethylation and promotes HIV-1 latency in primary CD4+ T cells (*40*). NFATC1, a calcium-responsive transcription factor, may inhibit LTR transcription by competing with NF-κB for overlapping promoter binding sites (*41*). RAD18, a DNA repair factor, has been reported to protect target cells against HIV-1 infection (*42*). The roles of PRAMEF16, *NDUFS5,* TAGLN2 and GBP4 in HIV-1 replication are unknown. GRN, HMOX1 and CIITA were top hits in the HIV-1 screen and already confirmed to exert antiretroviral activity (*13*). Although CEACAM3 restricts HIV-1 replication in primary CD4+ T cells (*13*) it exhibited higher MAGeCK scores in the SIVcpz*Ptt* screen (Fig. 4A, right).

**Fig. 4.**
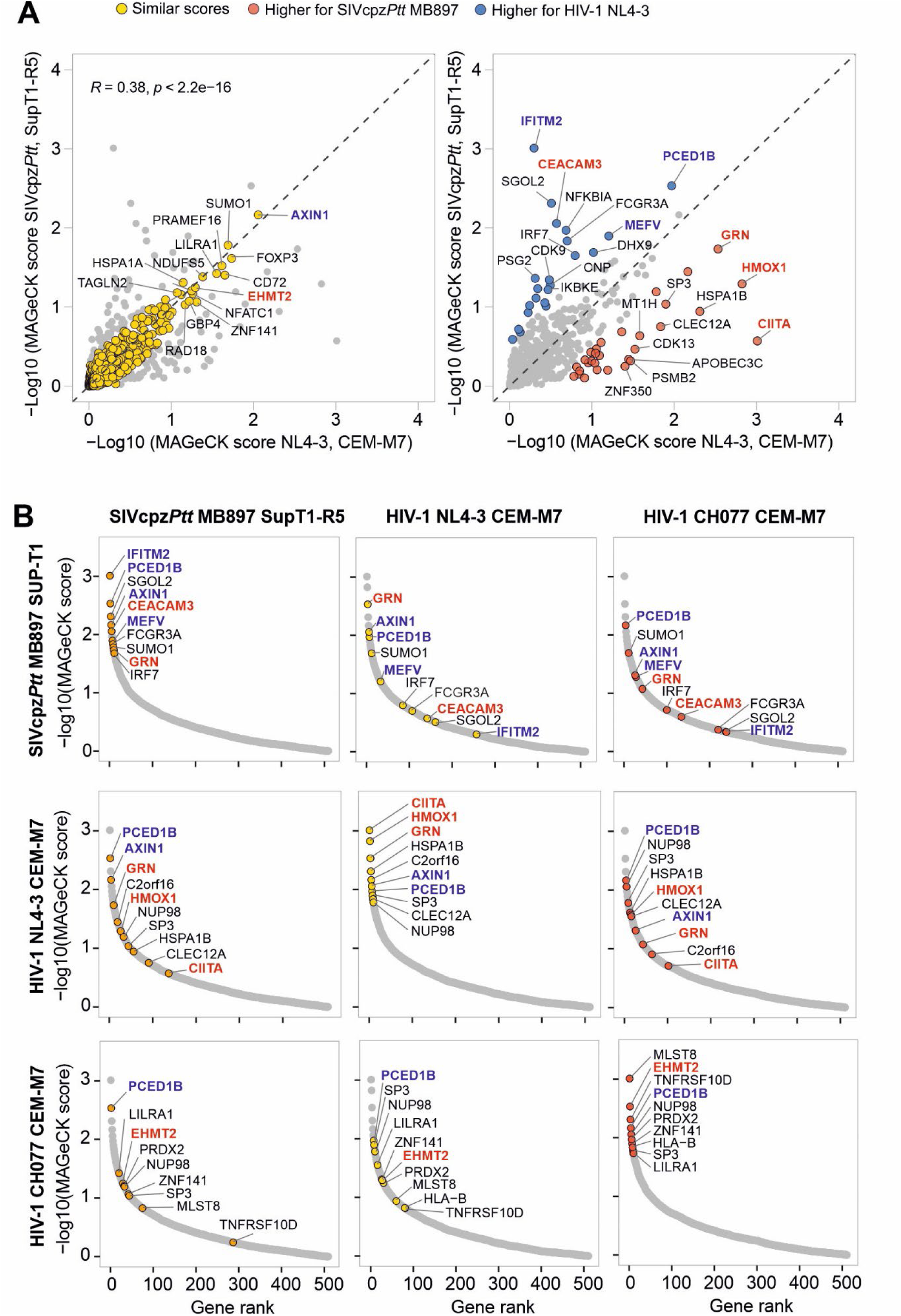
Overlaps and differences between sgRNAs enriched in SIVcpz- and HIV-1-driven screens. **(A)** MAGeCK scores obtained for each gene in the SIVcpz- and HIV-1-driven screens performed in SupT1-R5 and CEM-M7 Cas9 cells at 12 and 10 dpi, respectively. The left panel highlights genes with similar, and the right panel genes with different selection strengths by SIVcpz*Ptt* MB897 (red dots) or HIV-1 NL4-3 (blue dots). Factors selected for functional analyses are highlighted in blue and those analyzed in the previous HIV- 1 screen in red. (**B**) Top ten genes obtained in the SIVcpz and HIV-1 driven screens based on MAGeCK scores and their ranking in other screens at 12dpi.

To further define factors enhancing viral replication in a species- or strain-specific manner, we compared MAGeCK score-based rankings of the top 10 genes from the SIVcpz*Ptt* MB897, HIV-1 NL4-3, and CH077 screens (Fig. 4B). Fourteen genes consistently ranked among the top 50 hits in all screens: *PCED1B, SGOL2, AXIN1, CEACAM3, MEFV, FCGR3A, SUMO1, GRN, EHMT2, PRDX2, SP3, LILRA1, HSPA1B* and *HMOX1*. The corresponding cellular factors likely fall into several mechanistic clusters. Transcriptional repressors such as EHMT2, SP3 and SUMO1 may silence proviral gene expression through histone modification, SUMOylation, or NF-κB inhibition. Stress-response and antioxidant enzymes, like HMOX1, PRDX2, and HSPA1, likely dampen oxidative and inflammatory pathways that promotes viral transcription and replication. Immune regulatory molecules (MEFV, CEACAM3, FCGR3A, LILRA1 and GRN) may modulate innate immune sensing or facilitate clearance of infected cells. Nuclear transport and signaling factors such as SGOL2 and AXIN1 could alter nuclear pore function or transcriptional cofactor availability. Consistent with previous reports that different HIV-1 strains may yield distinct ISG hits (*43*, *44*), CIITA (class II major histocompatibility complex transactivator) ranked highly for NL4-3 but not for MB897 or CH077, whereas MLST8 (MTOR-associated protein, LST8 homolog) was specific to CH077 (Fig. 4B). Notably, overexpression of FCGR3A (Fc gamma receptor IIIa) has been previously shown to inhibit infectious HIV-1 production (*25*). Together, these results indicate that SIVcpz*Ptt* and HIV-1 select overlapping yet distinct sets of genes, with some shared targets differing in selection efficiency. We prioritized *IFITM2*, MEFV, *AXIN1*, and *PCED1B* for further analysis because sgRNAs targeting these genes were rapidly enriched and enhanced SIVcpzPtt MB897 replication in SupT1 Cas9 cells (Fig. 3). In addition, their roles in lentiviral restriction are poorly understood, and *IFITM2*, MEFV, and *PCED1B* sgRNAs were more efficiently selected in the SIVcpz screen (Fig. 4).

### IFITM2 efficiently restricts SIVcpz*Ptt* and some HIV-1 group M strains

IFITM3 is a well-established inhibitor of HIV-1 entry that blocks fusion of viral and host membranes (*45–47*). IFITM1 and IFITM2 are less well characterized and commonly considered weaker restriction factors. Unexpectedly, however, sgRNAs targeting *IFITM2* were enriched more efficiently than those targeting *IFITM3* (Fig. 5A). SIVcpz*Ptt* MB897 expressing an IFITM2-targeting sgRNA outcompeted an otherwise isogenic virus containing NT sgRNA (Fig. 3C), and produced higher levels of infectious virus in SupT1-R5 Cas9 cells (Fig. 3D). Overexpression of IFITM2 in HEK293T cells inhibited SIVcpz*Ptt* MB897 in a dose-dependent manner, while AXIN1, MEFV and PCED1B had little or no effect (Fig. S6). This agrees with findings that IFITMs incorporate into virions and reduce infectivity (*48*). To directly compare antiviral potency, HEK293T cells were cotransfected with IFITM1, IFITM2, or IFITM3 expression vectors and a panel of 15 different HIV-1 and SIVcpz IMCs. Western blots confirmed similar expression levels of different IFITMs (Fig. S7). Consistent with the results of the screen, IFITM2 showed the strongest antiviral activity, reducing infectious yields of HIV-1 M strains by an average of 85% and four SIVcpz*Ptt* IMCs by 92%, compared to 62% and 72% for IFITM1, and 70% and 75% for IFITM3, respectively (Fig. 5C).

**Fig. 5.**
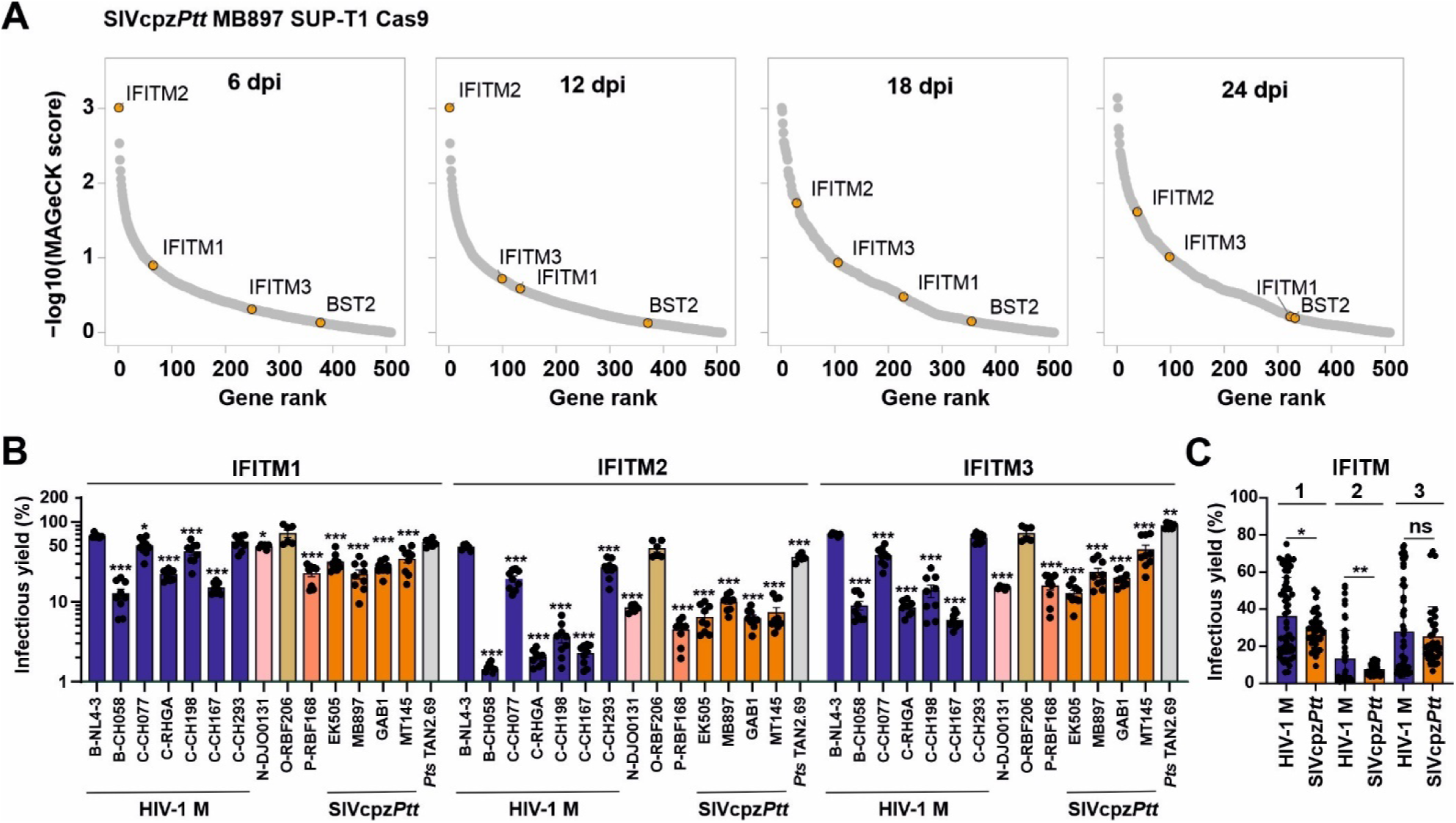
Enrichment of IFITM-targeting sgRNAs and antiviral activity of IFITMs. (**A**) Position of the three IFITM genes based on MAGeCK scores at the indicated days post-infection. BST-2 is shown for comparison. (**B**) HEK293T cells were transfected with IFITM expression vectors and the indicated HIV and SIV IMCs. Data show infectious virus yield relative to NT. P values are shown relative to those obtained for HIV-1 NL4-3. Dots represent data from three experiments each measured in duplicate or triplicate (mean ± SEM). (**C**) Average values (± SEM). obtained for all seven group M and the four SIVcpz*Ptt* strains. P values were determined using a two-tailed Student’s t-test with Welch’s correction. Significant differences are indicated as: *p < 0.05; **p < 0.01; ***p < 0.001; ns, not significant.

On average, SIVcpz*Ptt* strains were significantly more sensitive to IFITMs than HIV-1 IMCs. However, the sensitivity of HIV-1 group M strains varied substantially. Transmitted-founder HIV-1 IMCs CH058, RHGA, CH198 and CH167 were highly sensitive, while NL4-3, CH077 and CH293 were more resistant. HIV-1 N DJO0131 was also sensitive, whereas HIV-1 O RBF206 and SIVcpz*Pts* TAN2.69 were largely resistant against overexpression of IFITMs. Notably, SIVcpz*Ptt* MB897 was more sensitive to IFITM-mediated restriction than HIV-1 NL4-3 and CH077 used in prior screens (*13*), which agrees with stronger enrichment of IFITM2-targeting gRNAs by SIVcpz*Ptt* (Fig. 4). Altogether, the results show that IFITM2 overexpression in virus-producer cells inhibits SIVcpz*Ptt* more efficiently than IFITM1 and IFITM3. HIV-1 strains differ substantially in IFITM susceptibility, and the relative resistance of NL4-3 may explain why IFITM2’s antiviral activity has been underestimated in previous studies.

### Increased AXIN1 expression suppresses SIVcpz*Ptt* and HIV-1 replication

AXIN1 (Axis Inhibition Protein 1) emerged as a hit in both SIVcpz*Ptt*- and HIV-1-driven screens (Fig. 4), and sgRNAs targeting *AXIN1* strongly enhanced SIVcpz replication (Fig. 3). However, AXIN1 overexpression in HEK293T cells failed to inhibit SIVcpz*Ptt* MB897 and HIV-1 NL4-3 (Fig. S6). To further examine whether AXIN1 restricts viral replication in a cell type specific manner, we infected primary CD4+ T cells with SIVcpz*Ptt* MB897 and HIV-1 NL4-3 or CH077 in the presence of IWR-1, a tankyrase inhibitor that stabilizes AXIN1 (*49*). Treatment with IWR-1 significantly enhanced the levels of AXIN1 expression (Fig. 6A) without causing cytotoxic effects (Fig. 6B). IWR-1 treatment inhibited replication of SIVcpz*Ptt*, as well as both HIV-1 strains (Fig. 6C) and reduced infectious virus production by ∼50-60% (Fig. 6D) These findings indicate that artificially increased AXIN1 activity restricts SIVcpz*Ptt* and HIV-1 replication in primary T cells.

**Fig. 6.**
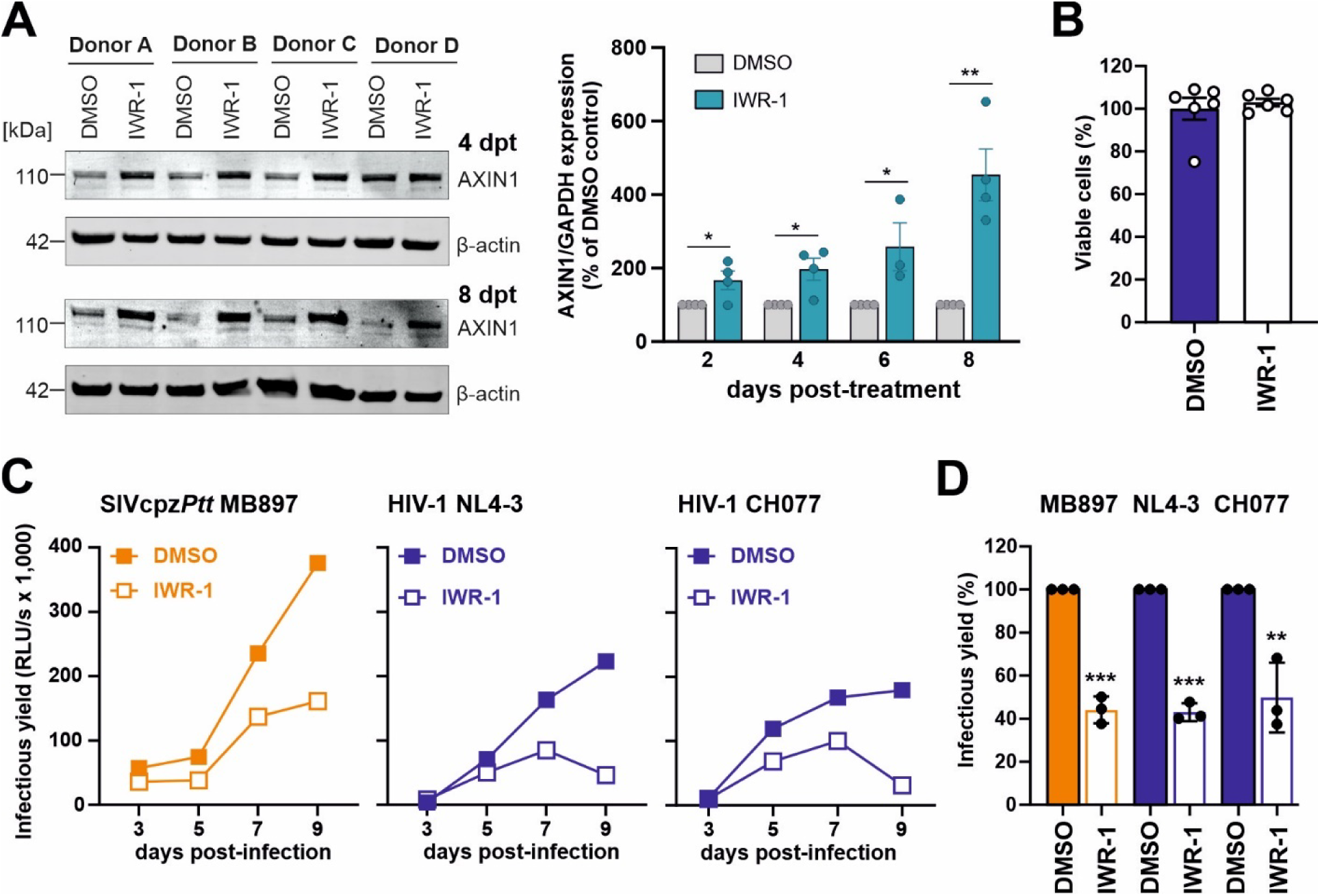
IWR-1 enhances AXIN1 expression and restricts viral replication. **(A**) Primary CD4⁺ T cells from healthy donors were treated every 2 days with 10 µM IWR-1 (dissolved in DMSO) or DMSO alone. Cells were harvested at the indicated time points for western blot analysis. The left panel shows AXIN1 expression in a representative donor at 4 and 8 days post-treatment. The right panel shows mean ± SEM percentages of AXIN1 expression in IWR-1–treated cells relative to DMSO controls (set to 100%). Statistical significance was determined using an unpaired t test (*p < 0.05; **p < 0.01). (**B**) Uninfected primary CD4⁺ T cells were treated with 10 µM IWR-1 or DMSO, and cell viability was assessed 2 days post-treatment using the CellTiter-Glo Luminescent Cell Viability Assay. (**C**) Primary CD4⁺ T cells were infected with SIVcpzPtt MB897, HIV-1 NL4-3, or CH077 and treated every 2 days with 10 µM IWR-1 or DMSO. Supernatants collected at 3, 5, 7, and 9 days post-infection were analyzed by TZM-bl assay. Data show representative replication kinetics from one of three donors, measured in two biological replicates. **(D)** Cumulative viral production from all three donors, normalized to DMSO controls (100%). P values were determined using a two-tailed Student’s t-test with Welch’s correction: **p < 0.01; ***p < 0.001.

### Host factors restricting SIVcpz*Ptt* but not HIV-1 in primary CD4^+^ T cells

To assess the potential *in vivo* relevance of factors identified by the SIVcpz*Ptt* TV screen, we confirmed expression of IFITM2, AXIN1, MEFV and PCED1B in primary CD4+ T cells (Fig. S8). For comparison, we also included BST-2. Although SIVcpz cannot antagonize BST-2 in human cells (*11*, *12*), it was not a significant hit in the screen since only one of three sgRNAs was enriched and increased infectious virus yields (Fig. 3). To further examine the antiviral activity of these factors, we used a previously established sgRNA/Cas9-based KO approach in primary CD4^+^ T cells (*13*). KO efficiently reduced protein levels of IFITM2, BST-2 and AXIN1, while depletion of MEFV and PCED1B was inefficient (Fig. S8). Reduced AXIN1, IFITM2 and BST-2 expression significantly enhanced SIVcpz*Ptt* MB897 replication, with the effect of BST-2 being stronger in the presence of IFN-β (Fig. 7A). To confirm inhibition of SIVcpz and examine susceptibility of HIV-1, we analyzed a second SIVcpz*Ptt* IMC (MT145) (*15*) and HIV-1 NL4-3. Partial KO of *IFITM2*, *AXIN1, MEFV* and (most strongly) *BST-2* significantly increased replication of SIVcpz*Ptt* MT145 but had no enhancing effect on HIV-1 NL4-3 (Fig. 7B). Altogether, the results revealed that endogenous levels of AXIN1, IFITM2, MEFV, and BST-2 restrict SIVcpz replication in primary human CD4+ T cells, whereas HIV-1 NL4-3 is resistant.

**Fig. 7.**
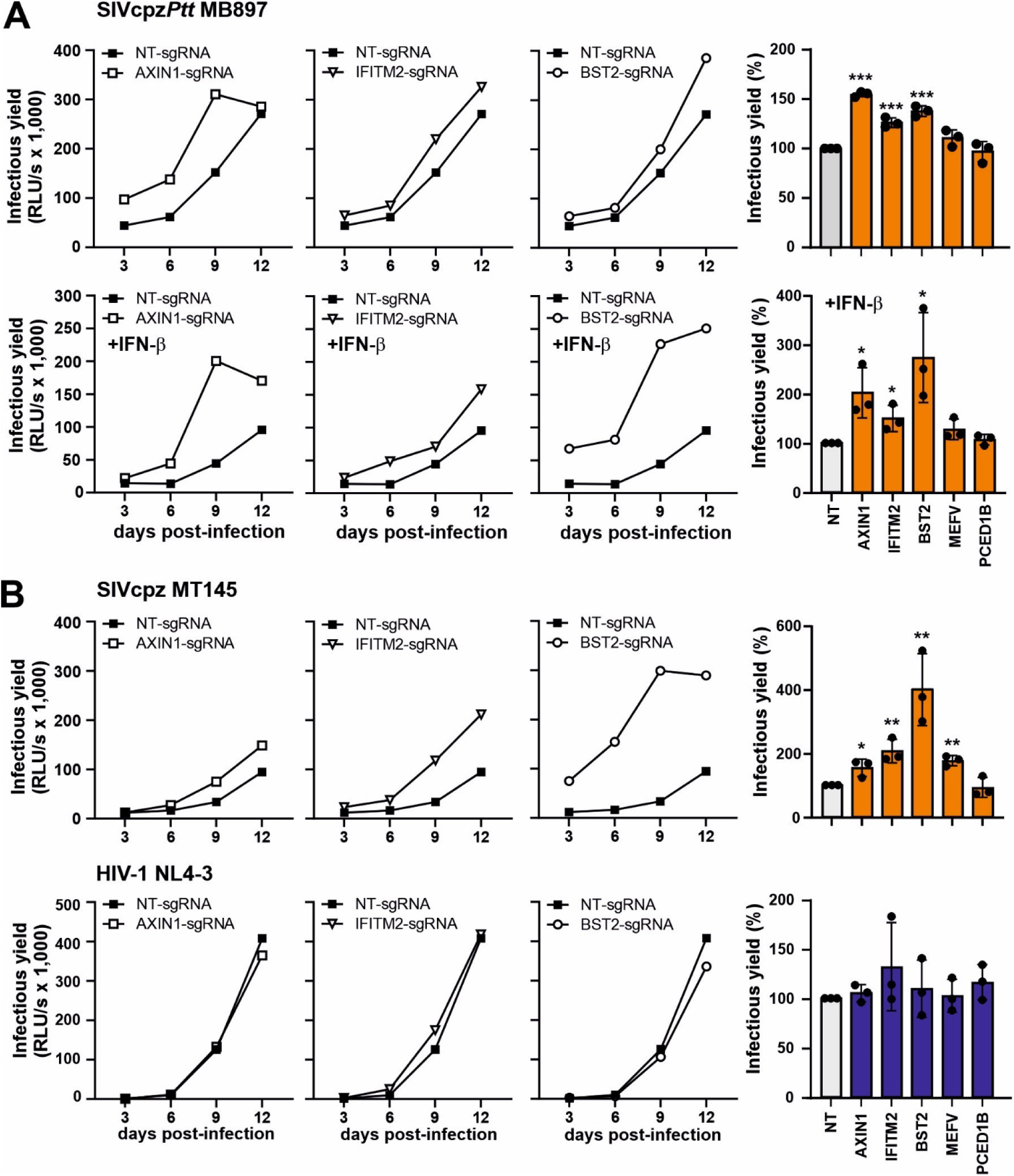
Effects of AXIN1, IFITM2 and tetherin KO on SIVcpz*Ptt* and HIV-1. (**A**) Primary CD4+ T cells were electroporated with AXIN1-, IFITM2-, BST-2-, or non-targeting (NT) sgRNAs followed by infection with SIVcpz*Ptt* MB897 in the absence (upper) or presence (lower) of IFN-β. Left panels: replication kinetics in cells from a representative donor. Each point represents the mean infectious virus yield from triplicate measurements. Right panels: cumulative infectious virus production by three donors, normalized to NT controls (100%). (**B**) Same as in panel A, except that cells were infected with SIVcpz*Ptt* MT145 or HIV-1 NL4-3. Only results in absence of IFN are shown since replication in presence of IFN-β was too low for meaningful analysis. P values were determined using a two-tailed Student’s t-test with Welch’s correction. Significant differences are indicated as: *p < 0.05; **p < 0.01; ***p < 0.001.

Conventional KO strategies, like the one above, are limited for studying antiviral factors because they alter cells before infection, potentially obscuring factors with distinct effects at different stages of viral replication. In addition, they may impact cell viability, especially since non-infected bystander cells are also affected. Furthermore, KO of MEFV and PCED1B was inefficient (Fig. S8). To overcome these issues, we transduced stimulated primary CD4⁺ T cells from four independent donors with a VSVg-pseudotyped lentiviral vector expressing Cas9 (Fig. 8A, S9). Two days later, these cell cultures were infected with SIVcpz*Ptt* MB897 constructs encoding sgRNAs against *IFITM2, AXIN1, MEFV, PCED1B*, or *BST-2*, or a non-targeting control. All five targeting sgRNAs enhanced replication of SIVcpz*Ptt* MB897 in both the absence and presence of IFN-β (Fig. 8B), and this enhancement was dependent on Cas9 expression (Fig. S10A). These results suggest that IFITM2, AXIN1, MEFV, PCED1B and BST-2 restrict SIVcpz replication in primary human T cells at both basal and IFN-inducible expression levels. In contrast, analogous HIV-1 NL4-3 and CH077 constructs carrying sgRNAs against *AXIN1, MEFV, PCED1B*, or *BST-2* did not show increased replication (Fig. 8C, S10B). On average, sgRNAs targeting all five factors identified by the SIVcpz-driven CRISPR screen significantly increased infectious SIVcpz*Ptt* MB897 production compared to the NT control, while none increased HIV-1 replication (Fig. 8D). Together, these results demonstrate that the TV approach identifies host factors that restrict SIVcpz*Ptt*, but not HIV-1 group M strains, in primary human CD4⁺ T cells.

**Fig. 8.**
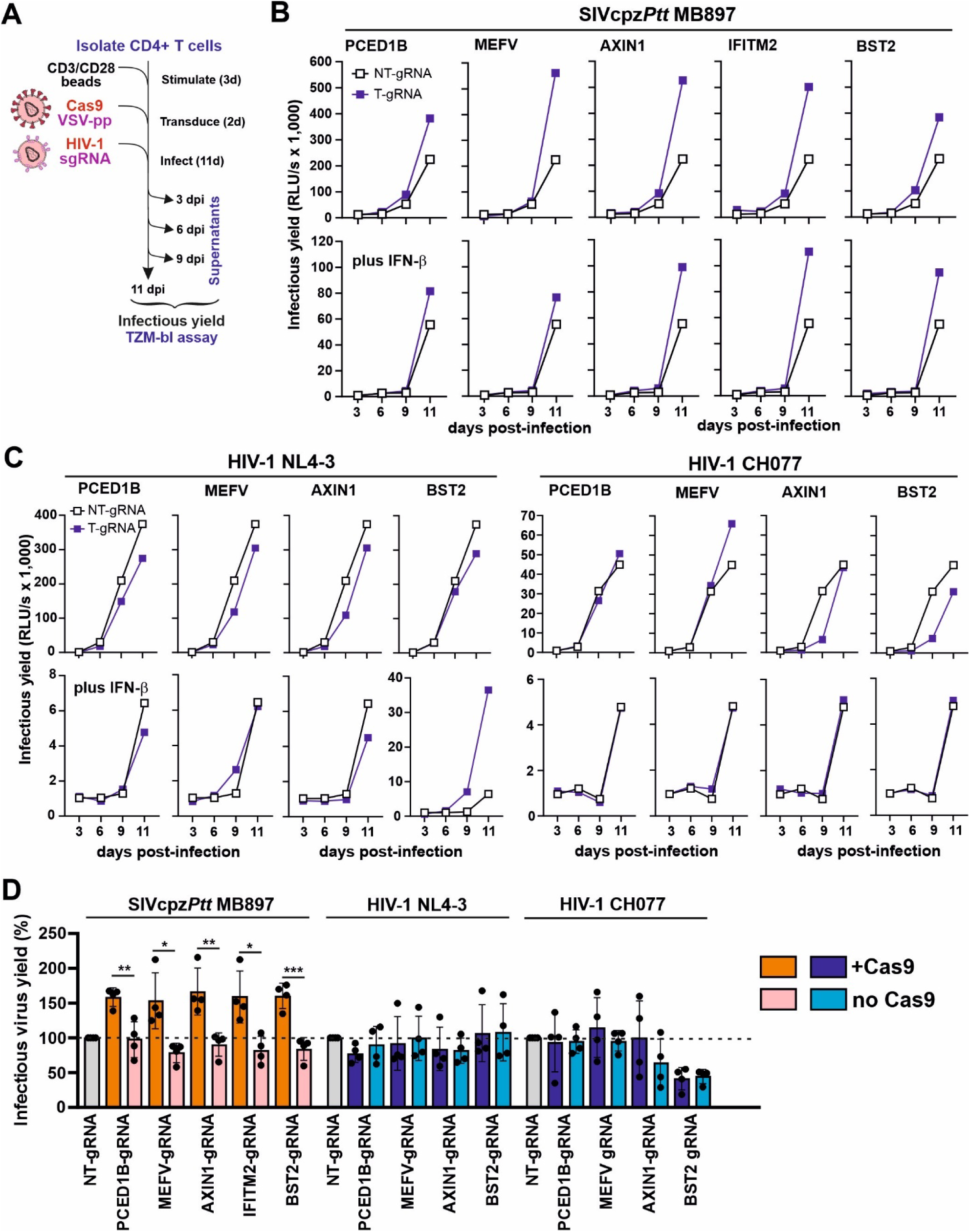
Effects of sgRNAs on SIVcpz and HIV-1 replication in Cas9-expressing CD4⁺ T cells. (**A**) Experimental schematic. (**B**) Replication of SIVcpzPtt MB897 constructs encoding sgRNAs targeting the indicated genes in Cas9-expressing primary CD4⁺ T cells. Infectious virus yields were measured by TZM-bl assay. (**C**) Replication of HIV-1 constructs the same sgRNAs as in (B). Panels (B) and (C) show data from the same representative donor as in (A). (**D**) Cumulative infectious yields from Cas9-expressing or control CD4⁺ T cells from four donors, normalized to the non-targeting (NT) sgRNA control (100%). Data represent means ± SEM from four independent experiments. P values were calculated using two-tailed Student’s t-tests with Welch’s correction. Significance: *p < 0.05; **p < 0.01; ***p < 0.001.

## DISCUSSION

Only one of at least four zoonotic transmissions of simian immunodeficiency viruses from chimpanzees (SIVcpz) or gorillas (SIVgor) gave rise to the AIDS pandemic (*8*, *9*). Identifying human host factors that restricted SIVcpz provides insights into the evolutionary barriers HIV-1 group M strains overcame to spread efficiently in humans. Using a replication-competent SIVcpz-driven CRISPR–Cas9 screen, we identified a variety of host genes including *IFITM2*, *AXIN1, MEFV* and *PCED1B*, that restrict SIVcpz*Ptt* MB897 replication in human SupT1 cells. Functional analyses confirmed that these factors inhibit SIVcpz*Ptt* but not HIV-1 replication in primary human CD4+ T cells. These findings reveal host defense mechanisms that likely shaped lentiviral evolution and the emergence of pandemic HIV-1.

IFITM2 emerged as a top hit in the SIVcpz*Ptt*-driven CRISPR screen (Fig. 4B). IFITMs are known to be incorporated into budding virions and reduce HIV-1 infectivity (*46*, *48*, *50*, *51*). However, it was unexpected that sgRNAs targeting IFITM2 were selected more efficiently than those targeting IFITM3 (Fig. 5A) since it is commonly thought that the antiviral potency ranks IFITM3 > IFITM2 > IFITM1 (*48*, *52*). Consistent with the results of the CRISPR screens, overexpression of IFITM2 reduced the infectious yields of diverse HIV-1 and SIVcpz IMCs more efficiently than IFITM3 (Fig. 5B, 5C). On average, SIVcpz*Ptt* IMCs were more susceptible to IFITM2 restriction than HIV-1 group M strains, but the sensitivity of the latter varied considerably. Notably, NL4-3 and CH077 used in our previous CRISPR screens were largely resistant, whereas some TF HIV-1 IMCs showed marked sensitivity to IFITM2 and IFITM3 restriction (Fig. 5B). This was unexpected, since it has been reported that TF HIV-1 strains are relatively resistant to IFITM-mediated restriction (*47*). Altogether, the relative antiviral potency of IFITMs is dependent on both the assay and the viral context. Most previous studies relied on IFITM-overexpressing cell lines, whereas our data show that IFITM2 restricts SIVcpz*Ptt* MB897 and MT145, but not HIV-1 NL4-3, in primary human T cells. These findings support a role of IFITM2 in innate defense against SIVcpz*Ptt* and suggests adaptation to the human host for efficient viral transmission. It has recently been reported that Nef counteracts HIV-1 restriction by IFITM2 and 3 (*53*) and it will be important to clarify whether Nef determines primate lentiviral sensitivity to IFITM proteins. Notably, SIVcpz*Pts* TAN2.69 from eastern chimpanzees was largely resistant to IFITM overexpression (Fig. 5B). Thus, differences in susceptibility to IFITMs do not explain why only SIVcpz*Ptt* has been detected in humans.

AXIN1, a central scaffold protein in the Wnt signaling pathway (*33*), was identified as another potent inhibitor of SIVcpz*Ptt* replication. AXIN1-targeting sgRNAs significantly increased SIVcpz*Ptt* MB897 replication in both SupT1 cells and primary human CD4+ T cells in a Cas9-dependent manner (Fig. 3D, Fig. 8). In contrast, overexpression of AXIN1 in HEK293T cells had little effect on infectious virus yields (Fig. S6), suggesting a context-dependent function that may be specific to T cells. Previous studies on HIV-1 reported that Wnt signaling inhibits viral replication and promotes viral latency in a ß-catenin dependent manner (*54*, *55*). AXIN1 is a negative regulator of the Wnt pathway (*33*). Thus, its identification as an effective inhibitor of SIVcpz*Ptt* was unexpected but agrees with recent data showing that AXIN1 boosts the antiviral response through IRF3 stabilization (*56*). AXIN1-targeting sgRNAs were enriched in the context of both SIVcpz*Ptt* and HIV-1 (Fig. 4) and artificially enhanced AXIN1 expression inhibited replication of both SIVcpz and HIV-1 (Fig. 6). In contrast, endogenous AXIN1 expression inhibited replication of SIVcpz*Ptt* but not HIV-1 in primary CD4+ T cells (Fig. 8). These findings identify AXIN1 as a T cell–specific antiviral factor for SIVcpz*Ptt* and suggest that reduced susceptibility to AXIN1-mediated inhibition facilitates efficient spread of HIV-1 in humans.

MEFV (Mediterranean fever gene) and PCED1B (PC-esterase domain containing 1B) were also identified as inhibitors of SIVcpz*Ptt* MB897 replication. In SupT1-R5 Cas9 cells, sgRNAs targeting these genes significantly enhanced viral replication (Fig. 3D), suggesting roles in the innate antiviral defense. MEFV encodes pyrin, a regulator of inflammasome activity and inflammatory signaling, that has been implicated in controlling human pathogens and in induction of autophagy and autoinflammatory diseases (*57–60*). Further studies are required to determine whether its antiviral activity involves inflammasome-mediated restriction or modulation of cellular stress responses during HIV-1 infection. PCED1B is less well characterized, but its predicted esterase activity suggests it may influence lipid metabolism or membrane composition, potentially impacting viral assembly or egress. Overexpression of MEFV or PCED1B in HEK293T cells did not reduce infectious viral yields (Fig. S6), suggesting that their antiviral effects may be cell-type specific. Indeed, MEFV and PCED1B targeting sgRNAs significantly enhanced replication of SIVcpz*Ptt* but not of HIV-1 in primary human CD4+ T cells (Fig. 8, S10). These findings identify MEFV and PCED1B as novel antiviral factors for SIVcpz*Ptt* and suggest that resistance to their antiviral effects may have contributed to the successful adaptation of HIV-1 to humans.

Acquisition of Vpu as an effective tetherin antagonist was a major adaptation of HIV-1 group M to the human host and most likely played a key role in its pandemic spread (*8*). Nevertheless, tetherin was not among the top hits in our SIVcpz*Ptt*-driven CRISPR screen (Fig. 3). In part, this may reflect its low expression in SupT1 cells and the virus’s preferential cell-to-cell transmission mode (*27*, *28*). However, it also indicates one limitation of our approach - its reliance on the efficiency and specificity of individual sgRNAs. Tetherin received a low MAGeCK score because only one of three targeting sgRNAs was enriched (Fig. 3C). Notably, sgRNA enhanced SIVcpz*Ptt* MB897 replication in both SupT1 cells and primary CD4+ T cells in a Cas9-dependent manner (Fig. 3D, 8B). In contrast, it did not enhance replication of HIV-1 NL4-3 and CH077, whose Vpu proteins efficiently antagonize human tetherin. A striking observation was the Cas9-independent enhancement of NL4-3 replication by the tetherin-targeting sgRNA in the presence of IFN-β (Fig. 8C). This increase was consistently observed in CD4⁺ T cells from all four donors (Fig. S10B) and warrants further investigation. Altogether, these findings confirm that human tetherin restricts SIVcpz but not HIV-1 and illustrate that inefficient sgRNAs may lead to underestimation or omission of bona fide restriction factors. However, continuous improvement of CRISPR-KO libraries, now achieving ∼70 % highly active sgRNAs, >95 % gene coverage, and <5 % off-target activity (*61*) will minimize this limitation. For most of our identified hits, at least two independent sgRNAs were significantly enriched during SIVcpz*Ptt* propagation (examples in Fig. 3C).

Since our screen is driven by viral replication fitness, with effects amplified over multiple rounds of replication, it is highly sensitive and robust. A major advance of the present study is that stable lentiviral delivery of Cas9 expression vectors into primary CD4+ T cells enabled direct testing of enriched sgRNAs in the main target cells of SIV and HIV-1. This system verified that endogenous expression of PCED1B, IFITM2, AXIN1, MEFV and BST-2 restricts SIVcpz*Ptt* MB897 but not HIV-1 NL4-3 and CH077 in primary CD4+ T cells. While further studies are required to assess the variability of diverse SIVcpz and HIV-1 strains in susceptibility to inhibition, these data provide proof-of-concept that the TV approach allows to identify antiviral mechanisms that HIV-1 M has cleared during adaptation to humans. As also noted in our previous study (*13*), most factors limiting viral replication fitness were not ISGs. Antiviral factors that are constitutively expressed may represent the under-investigated real first line of defense as they do not require viral replication and innate immune activation to exert protective effects.

The TV approach offers many prospects for further studies. Constitutive or conditional Cas9-expressing mouse models are already in routine use and the development of analogous non-human primate models is technically feasible (*62*). Thus, screening of TV-sgRNA libraries or verification of individual sgRNA in animal models for HIV/AIDS offers interesting perspectives. Notably, the TV approach not only identifies specific antiviral factors but also host genes affecting cellular pathways and general features that are not favorable for effective viral replication. Thus, genome-wide TV screens using different viral backbones, cell types, and experimental conditions should provide unprecedented deep new insights into the virus-host interplay.

## MATERIALS AND METHODS

### Replication-competent SIVcpz*Ptt* MB897 sgRNA constructs

Generation of the pCR-XL-TOPO-SIVcpz MB897 IMC has been previously reported (*15*). To generate replication-competent SIVcpz-MB897 containing an sgRNA cassette, the cassette was inserted between the *nef* gene and a duplicated TPI/U3 sequence, which usually overlaps *nef*. To prevent recombination, the TPI and overlapping *nef* regions were codon-optimized (Mut1). To reduce genome size, partial upstream U3 sequences (primarily encoding Nef and partially for transcription regulation) were removed (*22*), retaining TPI and either 89 bp (Mut2) or 111 bp (Mut3) of U3 upstream of the TATA box.

### Construction of traitor SIVcpz*Ptt* MB897 proviral vectors

DNA segments containing human codon optimized *nef*, sgRNA expression cassette, and truncated/duplicated 3’LTRs were synthesized by Twist Bioscience, PCR amplified using Forward primer: 5’-AAGGAG-TAGGGCCAGTCTCG-3’ and Reverse primer: 5’-CCTCTAGATGCATGCTCGAGC-3’, and cloned into the *NruI/NotI* digested pCR-XL-TOPO-SIVcpz MB897 backbone using Gibson Assembly. Products were transformed into XL2-blue MRF’ competent cells. All constructs were confirmed by Sanger sequencing in Microsynth Seqlab.

### Cloning of sgRNA library into traitor SIVcpz MB897 sgRNA cassette

To construct the traitor SIVcpz*Ptt* MB897 library, a DNA oligonucleotide library targeting 510 human genes (*13*) was synthesized by Twist Bioscience. The synthesized DNA oligonucleotides were amplified by polymerase chain reaction (PCR) using NEBNext® High-Fidelity 2×PCR Master Mix (NEB, M0541L) and purified using the Monarch PCR & DNA Cleanup Kit (NEB, T1030L). The traitor SIVcpz*Ptt* MB897 vector was digested with Esp3I (Thermo Fisher Scientific, FD0454), and the amplified sgRNA sequences were assembled into the vector backbone with the Gibson Assembly method. Ligation products were purified using the Monarch PCR & DNA Cleanup Kit (NEB, T1030L). 200 ng of the purified ligation products was electroporated into 25 μl of Endura Competent Cells (Lucigen, 60242). Approximately 2.5×10^6^ recombinants were obtained, which yielded 1500× library coverage. Bacteria were harvested, and the sgRNA library plasmids were extracted using the QIAGEN Plasmid Maxi Kit (Qiagen, 12165).

### Infection, kinetics and traitor virus enrichment

To start the replication kinetic, 5 million SupT1 CCR5 Cas9 cells were infected with the traitor SIVcpz*Ptt* MB897 library constructs with the EF-C fibril enhancer (*63*) at a final concentration of 10 μg/ml. On the next day, cells were washed three times with PBS and seeded in T25 flask at a cell density of 0.7 million/ml in the presence or absence of 100 U/ml IFN-β (R&D Systems, 8499-IF-010). Every two to three days, cells were sub-cultured until 12 days post infection. At 12 d p.i., supernatants containing viruses were concentrated by Amicon Ultra 15mL Filters 100 kDa, to infect fresh SupT1 CCR5 Cas9 cells as above, and cultured for another 12 days. Every 6 days, supernatants were collected and concentrated for further analysis. Kinetics were monitored by infecting TZM-bl cells.

### Detection of sgRNA cassette stability by reverse transcription PCR

To check the stability of the sgRNA cassette in the viral genome during passaging, viral RNA was isolated at different time points from supernatants with the QIAamp Viral RNA Mini Kit (Qiagen, 52906). Residual genomic DNA was digested and cDNA was synthesized using the PrimeScript RT Reagent Kit with gDNA Eraser (Takara, RR047A) according to the manufacturer’s instructions and specific primer 5’-GGGCAAGCCACTCCCTACC-3’. The cassette was amplified from the cDNA by KOD ONE DNA polymerase (TOYOBO, KMM-201NV) using Forward primer: 5’-GGATGGCCTGCAGTAAGG-GAC-3’, Reverse Primer: 5’-GGGCAAGCCACTCCCTACC-3’. PCR reactions were loaded onto a 1% agarose and ran at 140 V for 30 min.

### Viral infectivity

To determine infectious virus yield, 8,000 TZM-bl reporter cells/well were seeded in 96-well plates and infected with cell culture supernatants in triplicates on the following day. Three days post-infection, cells were lysed and β-galactosidase reporter gene expression was determined using the X-Gal Screen Kit (Applied Bioscience, T1027) according to the manufacturer’s instructions with an Orion microplate luminometer (Berthold).

### CellTiter-Glo Luminescent Cell Viability Assay

Cell viability was assessed using CellTiter-Glo Luminescent Cell Viability Assay (Promega G7571), following the manufacturer’s protocol. Briefly, two days after IWR-1 treatment, 0.3 million uninfected primary CD4+ T cells in 150 µl culture medium were mixed with 150 µl of CellTiter-Glo Reagent. From this mixture, 50 µl was transferred into fresh plates in triplicate. After a 10-minute incubation at room temperature, luminescence was measured using an Orion microplate luminometer

### Flow cytometry

∼200,000 cells were harvested, washed once with PBS and stained for 30 min at RT in the dark with eBioscience Fixable viability dye 780 (ThermoFisher Scientific, 65-0865-18) and anti-CD4 antibody (PerCP-Cy5.5, Biolegend #317428). Afterwards, cells were washed twice with PBS and permeabilized 20min with 200 μl BDCytofix/Cytoperm Fixation/Permeabilization Solution Kit (BD Biosciences, 554714) at RT. Cells were washed twice with 200 μl 1× Perm/Wash solution and stained 1 h at 4 °C with anti-capsid antibody (KC57-FITC, Beckman Coulter, 6604667). After washing twice with 1× Perm/Wash solution, cells were resuspended in PBS and measured with BD FACSCanto II Flow Cytometer (BD Biosciences).

### Next Generation sequencing

NGS was performed using the Illumina NextSeq2000 platform with 60 base-pair paired-end runs. Raw reads were demultiplexed, trimmed, groomed according to quality and aligned to the custom library sequences using the MAGeCK algorithm suite on the Galaxy platform. Individual read counts were determined and median-normalized to account for differences in library size and read count distributions. Individual sgRNAs targeting the same gene were aggregated, and a variance model based on a negative binomial distribution was applied to statistically assess differences between control (input) and experimental conditions (different days). Gene targets were ranked by MAGeCK based on their *p*-values via a modified robust ranking aggregation (RRA) algorithm (α-RRA) to identify significantly enriched genes. Overrepresented sgRNA sequences compared to the input control represent viruses that had a sgRNA targeting a gene, that restricts viral replication. Volcano plots, correlation analyses, dotplots and heatmaps were generated using R version 4.4.2, ggplot2 version 3.5.2.

### SDS-PAGE and Immunoblotting

Western blotting was performed as previously described (*64*). In brief, whole cell lysates were mixed with 4x Protein Sample Loading Buffer (LI-COR, at a final dilution of 1x) supplemented with 10% β-mercaptoethanol (Sigma Aldrich), heated at 95°C for 5 min, separated on NuPAGE 4±12% Bis-Tris Gels (Invitrogen) for 90 minutes at 100 V and blotted onto Immobilon-FL PVDF membranes (Merck Millipore). The transfer was performed at a constant voltage of 30 V for 30 minutes using semi-dry transfer system. For larger proteins (Cas9, EHMT2), transfer was performed at a constant Amperage 0.4 A for 2 hours using a wet transfer system. After the transfer, the membrane was blocked in 1 % Casein in PBS (Thermo Scientific).

### CRISPR/Cas9 KO in primary human CD4+ T cells

CD4⁺ T lymphocytes were isolated from healthy donors. Cells were stimulated for 3 days with IL-2 (10 ng/ml; Miltenyi Biotec, #130-097-745) and anti-CD3/CD28 beads (Gibco, #11132D). Cultures were maintained in RPMI-1640 medium supplemented with 20% FCS and IL-2 (10 ng/ml). One million stimulated primary CD4⁺ T cells were electroporated with a HiFi Cas9 Nuclease V3 (IDT)/gRNA complex (80 pmol Cas9 300 pmol gRNA; Lonza), using either a non-targeting or gene-specific sgRNA. Electroporation was performed using the Amaxa 4D-Nucleofector™ system with the Human Activated T Cell Nucleofector™ Kit (P3; Lonza, #V4XP-3032), pulse code EO115. Three days post-electroporation, one million cells per sample were infected with the indicated SIV or HIV-1 strains. From 3 to 12 days post-infection (dpi), supernatants were collected every 3 days and infectious virus production was quantified using the TZM-bl reporter cell assay.

### TV kinetics in primary human CD4+ T cells

CD4+ T lymphocytes were isolated and stimulated as described above. Stimulated primary human CD4+ T cells were infected with lentiviruses carrying Cas9. At three days post-infection, 0.5 million cells per sample were infected with the indicated SIV or HIV strains. From 3 to 11 days post-infection (dpi), supernatants were collected every 3 or 2 days and infectious virus yield was measured on TZM-bl reporter cells.

### Statistics

Statistical analyses were performed using GraphPad PRISM 10 (GraphPad Software). P-values were determined using a two-tailed Student’s t-test with Welch’s correction or Two-way ANOVA with Sidak’s multiple comparisons. Unless otherwise stated, data are shown as the mean of at least three independent experiments ± SEM. Significant differences are indicated as: *, p < 0.05; **, p < 0.01; ***, p < 0.001. Statistical parameters are specified in the figure legends.

## List of Supplementary Materials

Materials and Methods

Fig S1 to S10

## Acknowledgments

We thank Dré van der Merwe, Kerstin Regensburger, Jana-Romana Fischer, Daniela Krnavek and Martha Meyer for technical assistance and Christina Stürzel for help with the generation of proviral constructs. We dedicate this work to the memory of Helmut Blum, whose critical contributions to the deep sequencing analysis were invaluable. He passed away before the completion of this manuscript.

## Author contributions

Conceptualization: FK, KMJS

Methodology: QX, QW, SN, GG, AB, MV, DK

Investigation: QX, QW, SN, AB, SK, AG, HB

Resources: DG, GW

Formal analysis: SK, AG, HB, QW, GG

Visualization: QX, QW, SN, GG, SK, AG, HB, FK

Writing – original draft: FK

Writing – review & editing: All authors

Supervision: FK, KMJS

Project administration: FK, KMJS.

## Data and materials availability

Further information and requests for resources and reagents should be directed to and will be fulfilled by FK. All data are available in the main text or the supplementary materials.”

## Supplementary Materials

## Supplementary Text

### Materials and Methods

#### Cell culture

All cells were cultured at 37 °C in a 5% CO_2_ atmosphere. Human embryonic kidney 293T cells purchased from American Type Culture Collection (ATCC: CRL3216) were cultivated in Dulbecco’s Modified Eagle Medium (DMEM, Gibco) supplemented with 10% (v/v) heat-inactivated fetal bovine serum (FBS, Gibco), 2 mM L-glutamine (PANBiotech), 100 μg/ml streptomycin (PANBiotech) and 100 U/ml penicillin (PANBiotech). TZM-bl cells were provided and authenticated by the NIH AIDS Reagent Program, Division of AIDS, NIAID, NIH from Dr. John C. Kappes, Dr. Xiaoyun Wu and Tranzyme Inc. THP-1 (ATCC, TIB-202), U-937 (ATCC, CRL-1593.2), PM1 (NIH AIDS Reagent Program, ARP-3038), Jurkat T4 (CD4^+^ human leukemia T cells) (ATCC, TIB-152), SupT1 CCR5 high Cas9 (*13*), and THP-1 Cas9 cells were cultured in Roswell Park Memorial Institute (RPMI) 1640 medium supplemented with 10%(v/v) heat-inactivated fetal bovine serum, 2 mM L-glutamine, 100 µg/ml streptomycin, 100 U/ml penicillin. For selection and maintenance of transgene expression, 0.3 μg/ml puromycin and 10 μg/ml blasticidin S HCl were added to the SupT1 CCR5 high Cas9 culture medium to maintain CCR5 and Cas9 expression, respectively. THP-1 Cas9 cells were maintained with 10 μg/ml blasticidin S HCl to retain Cas9 expression.

#### Phylogenetic analyses

Evolutionary analyses were conducted using NGPhylogeny (https://doi.org/10.1093/nar/gkz303, https://doi.org/10.1093/nar/gkae268). Gag amino acid sequences (SIVcpz*Ptt* MB897.2: EF535994, HIV-1 M-B CH058.tf: JN944907, HIV-1 M-B CH077.tf: JN944941, HIV-1 M-B RHGA.cc: KC312535, HIV-1 M-B NL4-3 X4: AF003887, HIV-1 M-C CH198.tf: KC156130, HIV-1 M-C CH167.cc: KC156213, HIV-1 M-C CH293.cc: KC156216, HIV-1 N DJO0131: AY532635, SIVcpz*Ptt* EK505: DQ373065, SIVcpz*Ptt* MT145: DQ373066, SIVcpz*Ptt* Gab1: X52154, SIVcpz*Pts* Tan2.69 DQ374657, SIVgor CP684: FJ424871, HIV-1 N 04CM-1015-04: DQ017382, HIV-1 N YBF30: AJ006022, HIV-1 O MVP5180: L20571, HIV-1 O ANT70: L20587, HIV-1 O RBF206: KY112585, HIV-1 P 06CMU14788: HQ179987, HIV-1 P RBF168: GU111555) were obtained from the NCBI database and aligned using MAFFT. A phylogenetic tree based on sequence similarity is shown. Scale bar: 0.1 amino acid replacements per site. The phylogenetic tree showing distance-based relationship interference data based on nucleotide sequences was generated using the NGPhylogeny.fr FastME tool (https://ngphylogeny.fr/) and was visualized using iTOL (https://itol.embl.de/). The tree is drawn to scale, with branch lengths indicating the number of substitutions per site.

#### Transfection and production of viral stocks

HEK293T cells were transiently transfected using PEI 25K (Polysciences, 23966-100) at a ratio of 2 μg of PEI per 1 μg of DNA and the medium was replaced 4-6 h post-transfection. To test antiviral effects of potential restriction factors, pcDNA-based expression constructs were cotransfected with the proviral constructs. In titration experiments, empty vector was used to keep the total DNA amount constant. The transfected cells were incubated for 4-6 h before the medium was replaced by fresh DMEM with 2% FCS. To generate virus stocks, one day before transfection, 20 mio cells were seeded in 15 cm dishes in 20 ml medium to obtain a confluence of 80-90% at the time of transfection. For transfection, 40 μg of DNA was mixed with 80 μg PEI 25K, incubated 15 min at RT and added dropwise to the cells. 48 h post-transfection, the virus was harvested, centrifuged 5 min at 2000 rpm and concentrated 40 times using Amicon Ultra 15mL Filters 100 kDa (Merck, UFC910096). The concentrated viruses were aliquoted and stored at −80 °C.

#### Viral RNA preparation for sequencing

Viral RNA was isolated from concentrated supernatants collected from the traitor SIVcpz*Ptt* MB897 library infected cells at indicated time points using the Viral RNA Mini Kit (Qiagen) according to the manufacturer’s instructions. Genomic DNA was digested and cDNA was synthetized using the PrimeScript RT Reagent Kit with gDNA Eraser (Takara #RR047A) according to the manufacturer’s instructions and specific primer 5’-TAAAAAGTGGCTAGCGATCGC-3’. 12 of cDNA reactions for 1 sample were purified using the Monarch PCR Purification Kit (NEB, T1030L) and eluted in 20 μl ddH_2_O. The sgRNA cassette was amplified using the NEBNext® High-Fidelity 2× PCR Master Mix (NEB) and primers including Illumina adapters and 8-nt barcodes to allow Next-Generation Sequencing analysis. PCR reactions were purified using the Monarch PCR & DNA Cleanup Kit (NEB, T1030L) and eluted in 20 μl ddH_2_O. Next-generation sequencing NGS was performed using the Illumina NextSeq2000 platform with 60 base-pair paired-end runs. Raw reads were demultiplexed on the Galaxy version 23.0 platform, forward and reversed reads were merged with SeqPrep 0.2.2. Merged reads were trimmed by Cutadapt 4.4 then aligned to the custom library sequences using the MAGeCK algorithm suite (Version 0.5.9.2.4). Individual read counts are determined and median-normalized for the effect of library sizes and read count distributions. Individual sgRNAs targeting the same gene are summarized, and a variance model calculated using a negative binomial model to statistically assess the difference between control (input) and the conditions (different days). Targets are ranked by MAGeCK according to their p-value via a modified robust ranking aggregation (RRA) algorithm ( α-RRA) to identify enriched genes. Overrepresented sgRNA sequences compared to the input control represent viruses carrying an sgRNA targeting a gene, that restricts viral replication. Scatter plots were generated using R version 4.2.3, ggplot2 version 3.4.3, and smplot2 version 0.2.4.

#### Supernatants and whole cell lysates

To determine expression of cellular and viral proteins, cells were washed in PBS and subsequently lysed in Western blot lysis buffer (150 mM NaCl, 50 mM HEPES, 5mM EDTA, 0.1% NP40, 500 μM Na_3_VO_4_, 500 μM NaF, pH 7.5) or radioimmunoprecipitation assay (RIPA) buffer (50 mM Tris-HCl; pH 7.4, 150 mM NaCl, 1% (v/v) NP-40, 0.5% (w/v) deoxycholic acid (DOC), 0.1% (w/v SDS) supplemented with protease inhibitor (Roche, 1:500). After 5 min of incubation on ice, samples were centrifuged (4°C, 20 min, 14.000 rpm) to remove cell debris. The supernatant was transferred to a fresh tube, the protein concentration was measured with Pierce Rapid Gold BCA Protein Assay Kit (Thermofisher) and adjusted using Western blot lysis buffer. Supernatants were centrifuged on top of a 20% sucrose layer in at 21,000 g for 2 hours. The viral pellet was then lysed in Western blot lysis buffer with 4x Protein Sample Loading Buffer (LICOR) supplemented with 10% β-mercaptoethanol (Sigma Aldrich) and heated at 95°C for 5 min.

---

**Fig. S1.**
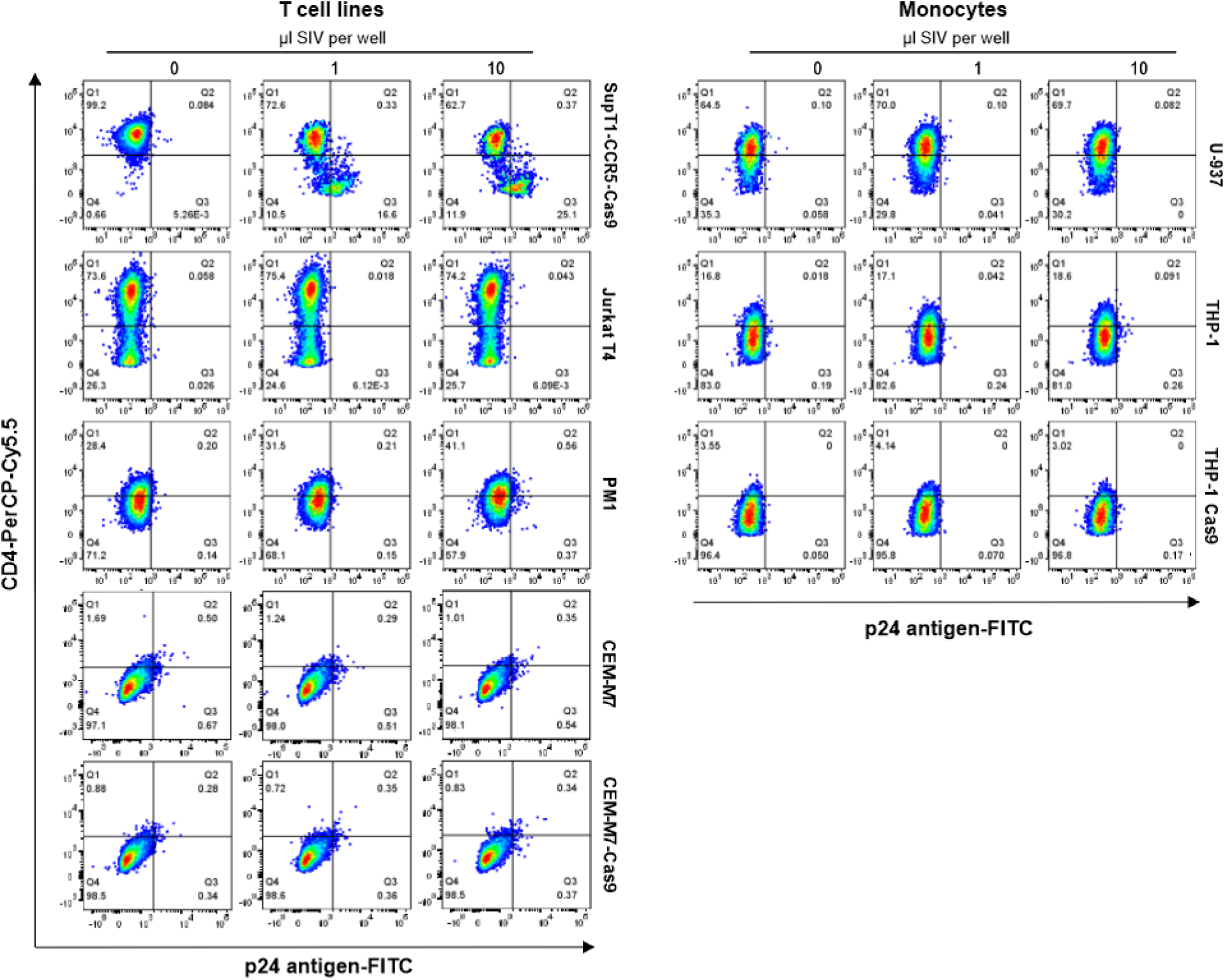
Susceptibility of human cell lines to SIVcpz*Ptt* MB897 infection. Flow cytometry analysis of CD4 and p24 capsid antigen expression levels in the indicated human cell lines after exposure to SIVcpz*Ptt* MB897.

**Fig. S2.**
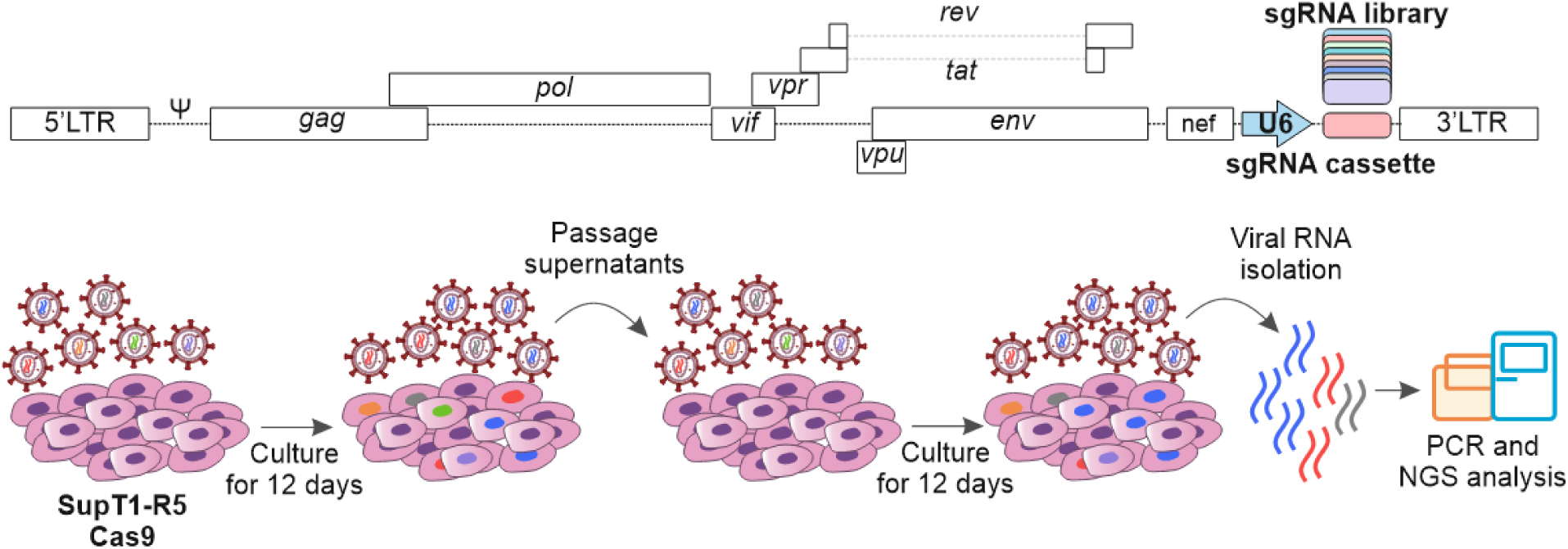
Schematic of proviral SIVcpz*Ptt* MB897 TV constructs and cell culture passaging. Proviral SIVcpz*Ptt* MB897 constructs are engineered to contain the sgRNAs expression cassette between the *nef* gene and the 3’LTR. To produce virus stocks, HEK293T cells are transfected with the viral libraries expressing various sgRNAs. The resulting swarms of MB897-sgRNA viruses are cultured for 12 days in Cas9-expressing cells in the presence or absence of IFN-β. Thereafter, the culture supernatant is used to initiate a 2^nd^ round of culture. Cells and viral supernatants are harvested every three days and the frequencies of sgRNAs are determined by next-generation sequencing. Note that the U6-sgRNA-scaffold region is not to scale.

**Fig. S3.**
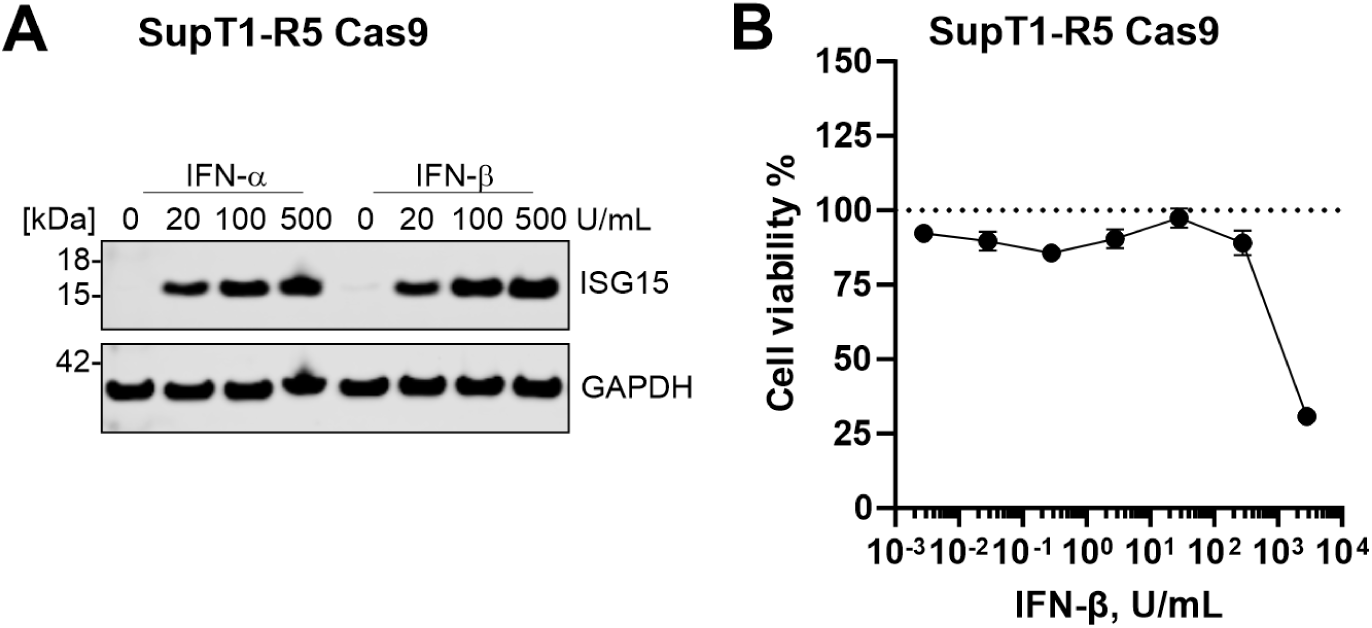
Responsiveness of SupT1-CCR5 Cas9 to type I IFNs. **(A)** Representative Western blot of ISG15 and GAPDH expression levels in SupT1-CCR5-Cas9 cells treated with the indicated doses of type I IFNs. **(B)** Effect of IFN-ß on the viability of SupT1-CCR5-Cas9 cells at 12 days post treatment. Shown are mean values (±SEM) from triplicate measurements.

**Fig. S4.**
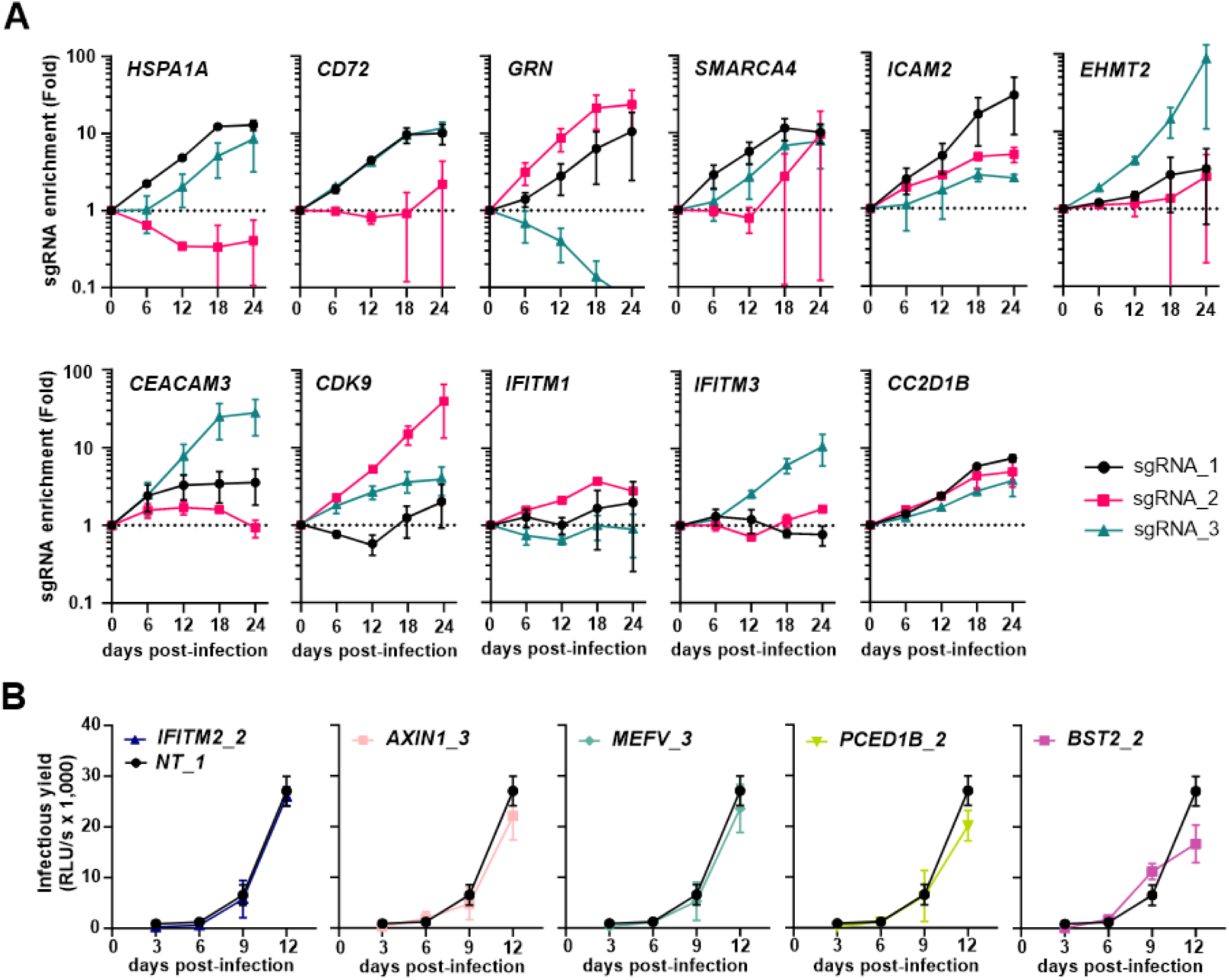
Enrichment of specific sgRNAs and enhancement of viral replication fitness. (**A**) Read-counts relative to input virus from the MAGeCK analysis showing the enrichment of sgRNAs targeting the indicated cellular genes over time. (**B**) SupT1-R5 cells were infected with SIVcpz*Ptt* MB897 constructs expressing the indicated targeting or a non-targeting control sgRNA. Infectious virus yield was measured using TZM-bl repoter cell. Symbols represent the mean of three independent experiments ±SEM.

**Fig. S5.**
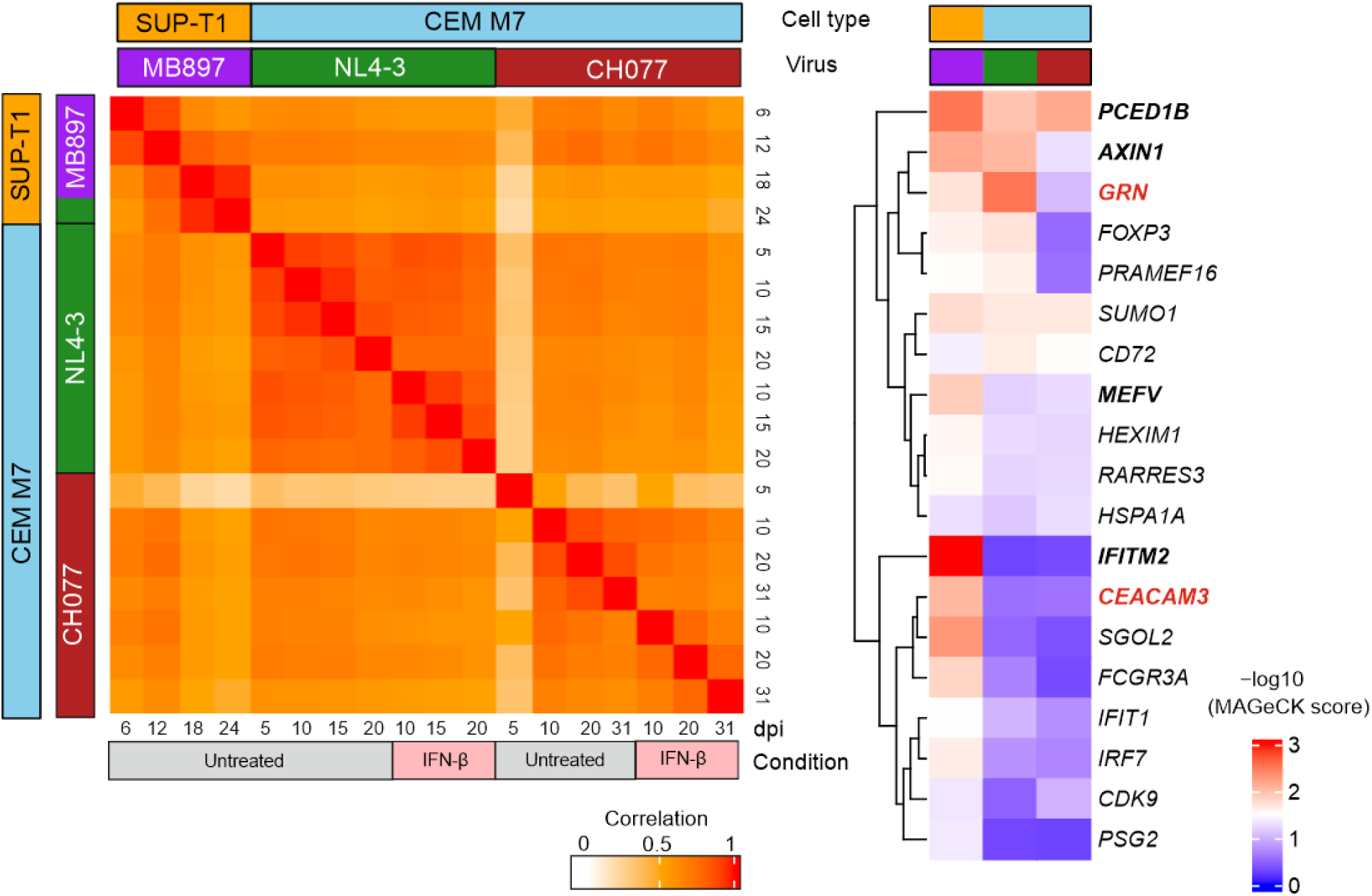
Comparison between SIVcpz*Ptt* MB897 and HIV-1 screens. (Left) Correlation of MAGeCK scores across all genes between the different HIV-1- and SIVcpz-driven screens. (Right) MAGeCK scores of all factors from the SIVcpz screen with log2 (fold-change) > 0 and p-value < 0.05 at 12 days post-infection, displayed across all screens.

**Fig. S6.**
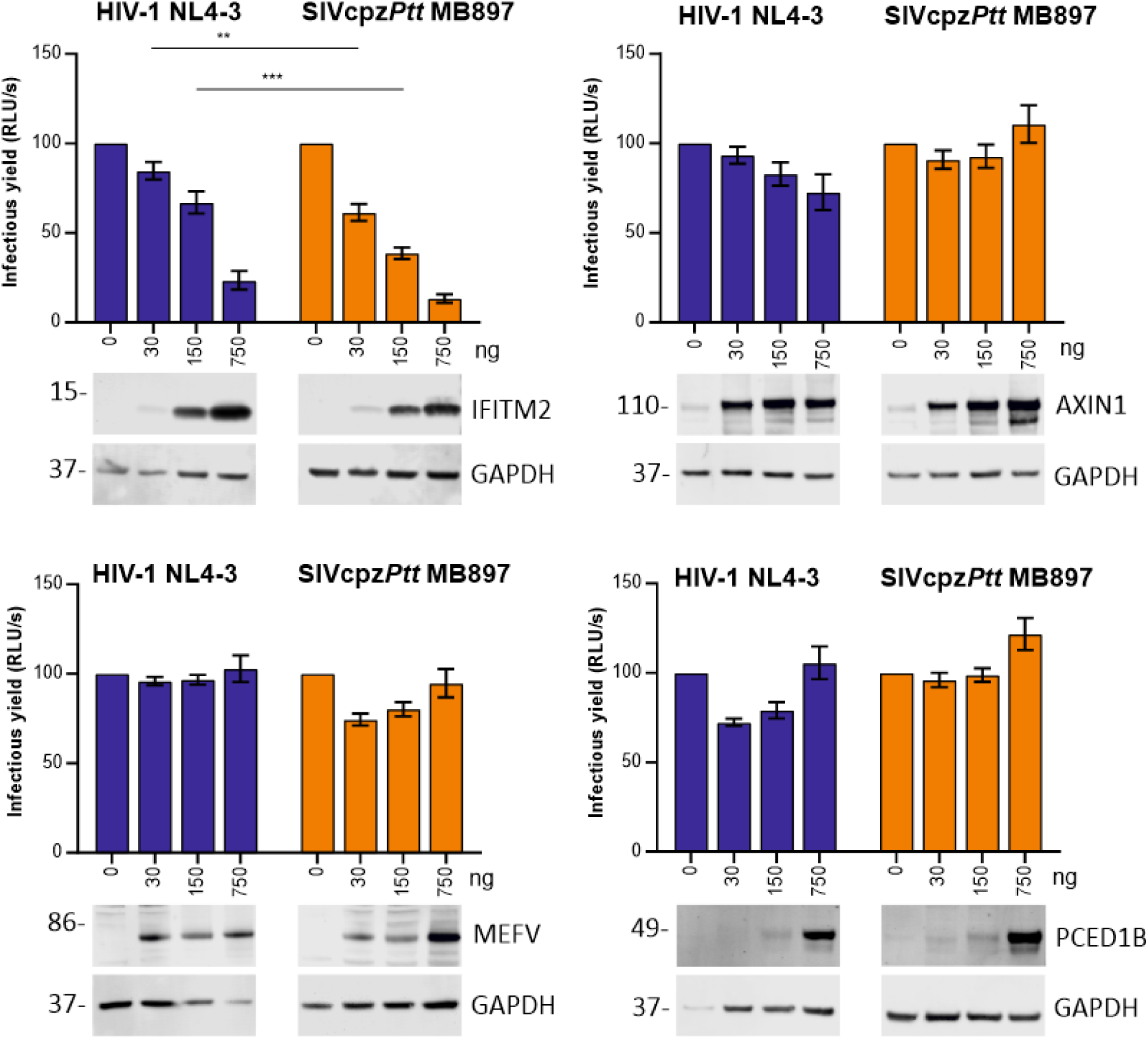
Effects of top hits overexpression in HIV-1 NL4-3 and SIVcpz MB897 production. HEK293T cells were co-transfected with increasing amounts of indicated expression vectors together with HIV-1 NL4-3 or SIVcpz MB897.2 proviral constructs. Supernatants were harvested at 2 days post-transfection, and infectious virus yields were measured on TZM-bl reporter cell. Shown are mean values (±SEM) from four independent experiments.

**Fig. S7.**
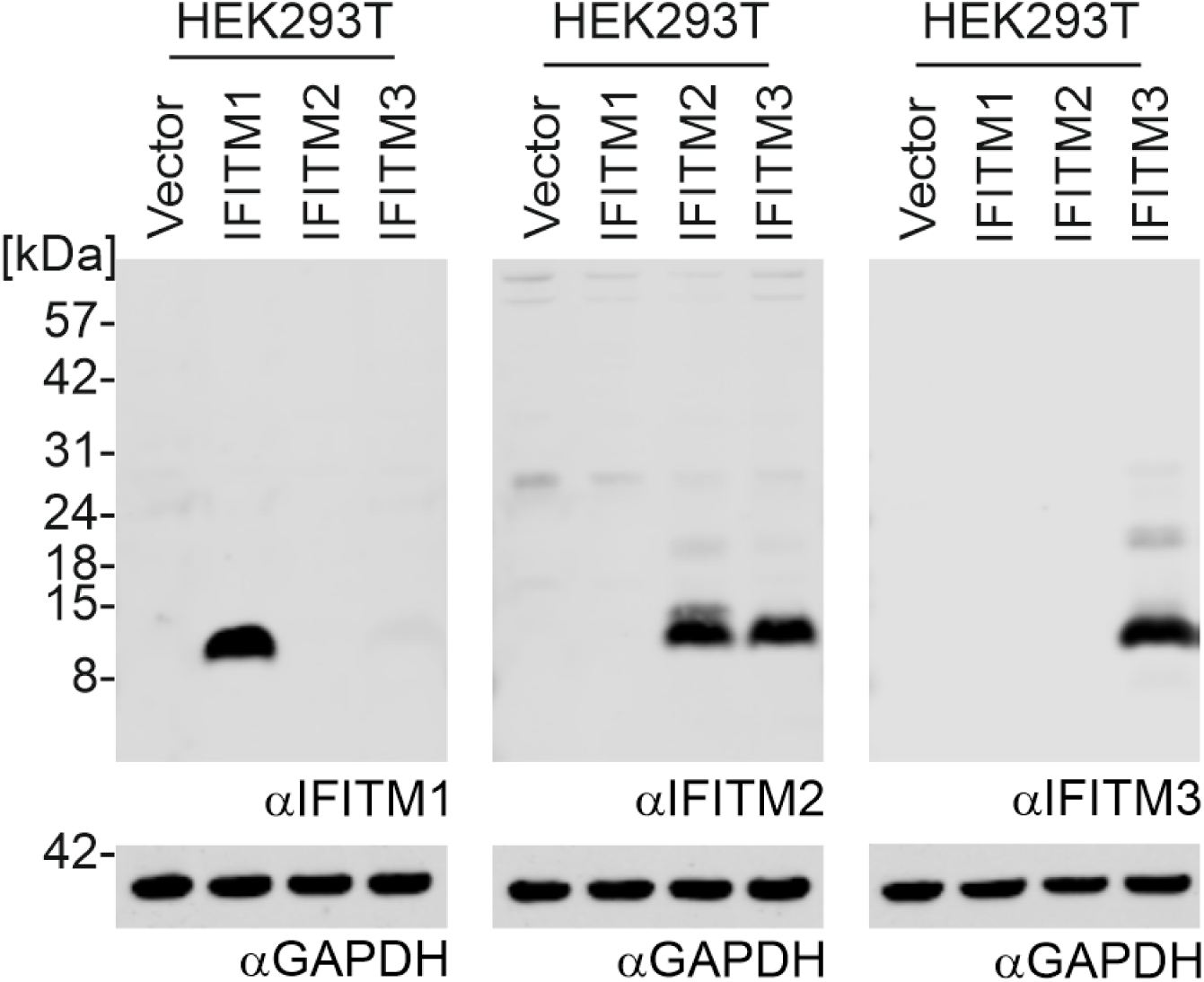
Expression of exogenous IFITMs proteins. HEK293T cells were co-transfected with IFITM expression vectors together with different IMCs. Whole cell lysates were harvested at 2 days post-transfection for western blot.

**Fig. S8.**
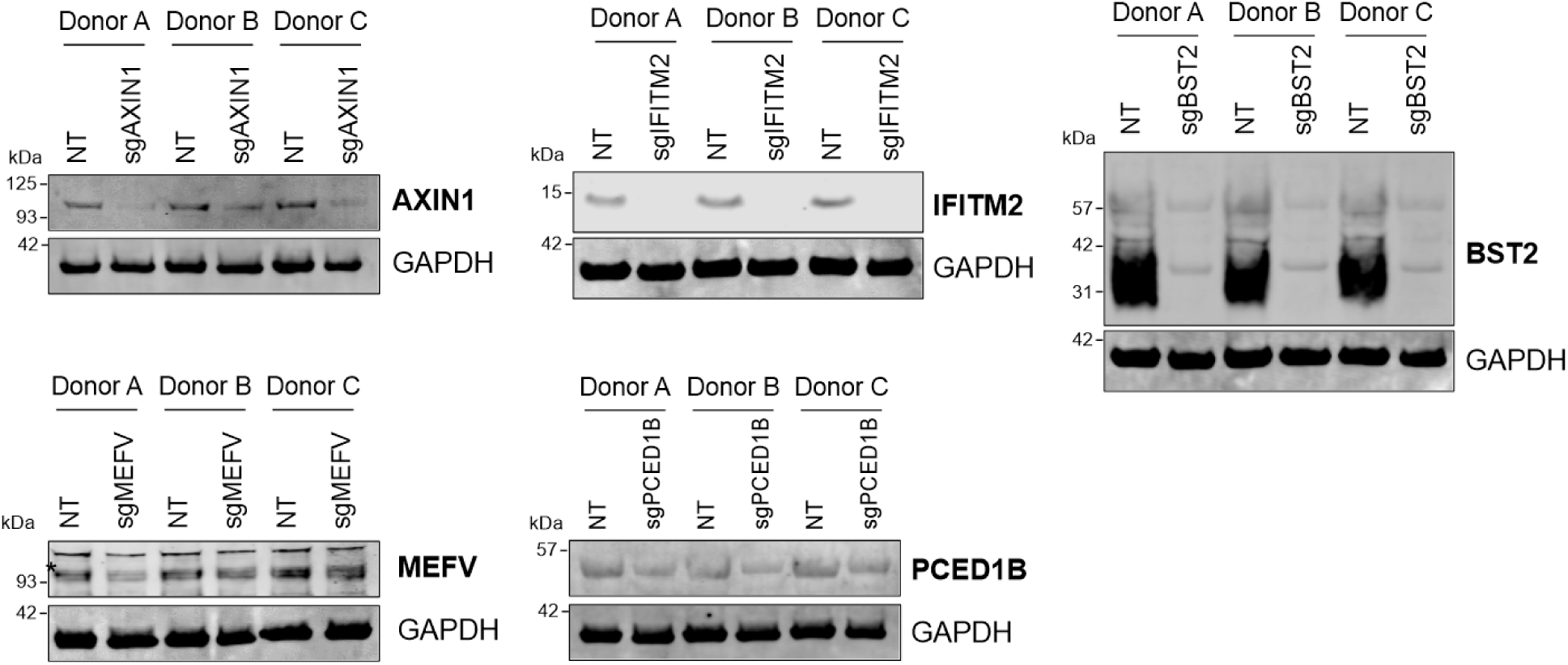
Knocking out efficiency in primary CD4+ T cells. Primary CD4+ T cells were electroporated with sgRNA targeting indicated genes or a non-targeting control. Whole cell lysates were harvested at 3 days post-electroporation for western blot.

**Fig. S9.**
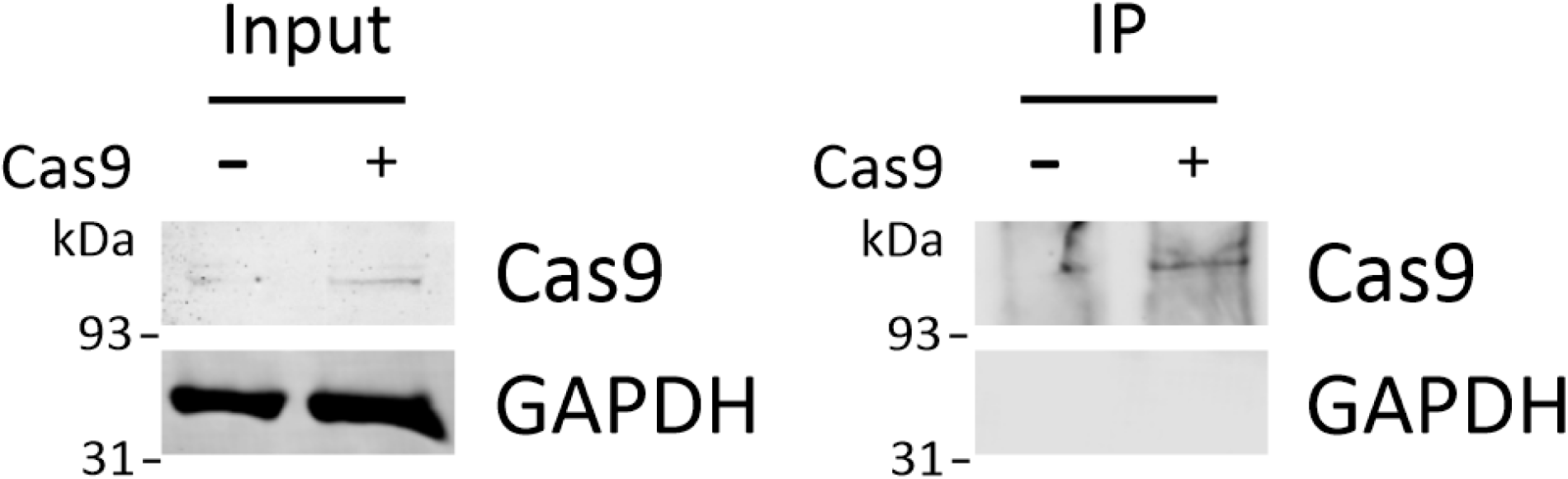
Verification of Cas9 expression in CD4+ T cells. Left: primary CD4+ T cells were transduced by a VSVg-pseudo-typed lentiviral vector expressing Cas9 or left untreated, harvested 2 days later, and analyzed by western blot. Right: to increase Cas9 signal intensity and ensure specificity, 1000 µg of protein lysates were subjected to immunoprecipitation using a Cas9 antibody.

**Fig. S10.**
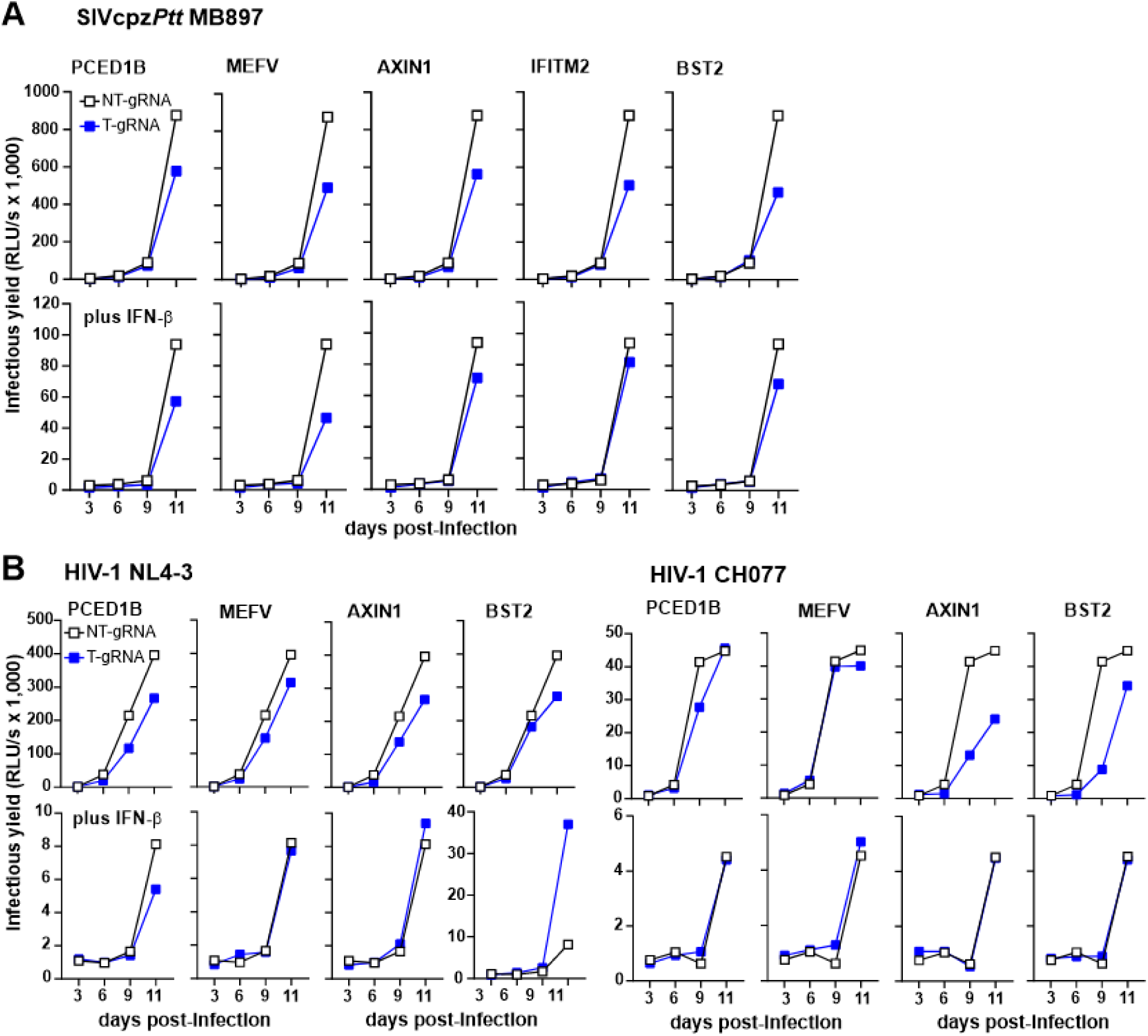
Effect of sgRNAs on SIVcpz and HIV-1 replication in primary human CD4+ T cells in the absence of Cas9. (**A**) Replication of SIVcpz*Ptt* MB897 sgRNA constructs targeting the indicated genes in primary human CD4+ T cells. Infectious virus yield was quantified by TZM-bl assay. (**B**) Replication of HIV-1 sgRNA constructs expressing the same sgRNAs as SIVcpz*Ptt* MB897.

## REFERENCES

1. J. G. N. Barreat, A. Katzourakis, A billion years arms-race between viruses, virophages and eukaryotes. eLife 12 (2023).

2. S. F. Kluge, D. Sauter, F. Kirchhoff, SnapShot: antiviral restriction factors. Cell 163, 774–774.e1 (2015).

3. M. H. Malim, P. D. Bieniasz, HIV Restriction Factors and Mechanisms of Evasion. Cold Spring Harbor Perspectives in Medicine 2, a006940–a006940 (2012).

4. M. Colomer-Lluch, A. Ruiz, A. Moris, J. G. Prado, Restriction Factors: From Intrinsic Viral Restriction to Shaping Cellular Immunity Against HIV-1. Front Immunol 9, 2876 (2018).

5. F. Kirchhoff, Immune evasion and counteraction of restriction factors by HIV-1 and other primate lentiviruses. Cell Host Microbe 8, 55–67 (2010).

6. L. Bergantz, F. Subra, E. Deprez, O. Delelis, C. Richetta, Interplay between Intrinsic and Innate Immunity during HIV Infection. Cells 8, 922 (2019).

7. M. S. Diamond, T.-D. Kanneganti, Innate immunity: the first line of defense against SARS-CoV-2. Nat Immunol 23, 165–176 (2022).

8. D. Sauter, F. Kirchhoff, Key Viral Adaptations Preceding the AIDS Pandemic. Cell Host & Microbe 25, 27–38 (2019).

9. P. M. Sharp, B. H. Hahn, Origins of HIV and the AIDS pandemic. Cold Spring Harbor perspectives in medicine 1, a006841 (2011).

10. F. Gao, E. Balles, D. L. Robertson, Y. Chen, C. M. Rodenburg, S. F. Michael, L. B. Cummins, L. O. Arthur, M. Peeters, G. M. Shaw, P. M. Sharp, B. H. Hahn, Origin of HIV-1 in the chimpanzee Pan troglodytes troglodytes. Nature 397, 436–441 (1999).

11. D. Sauter, M. Schindler, A. Specht, W. N. Landford, J. Münch, K.-A. Kim, J. Votteler, U. Schubert, F. Bibollet-Ruche, B. F. Keele, J. Takehisa, Y. Ogando, C. Ochsenbauer, J. C. Kappes, A. Ayouba, M. Peeters, G. H. Learn, G. Shaw, P. M. Sharp, P. Bieniasz, B. H. Hahn, T. Hatziioannou, F. Kirchhoff, Tetherin-Driven Adaptation of Vpu and Nef Function and the Evolution of Pandemic and Nonpandemic HIV-1 Strains. Cell Host & Microbe 6, 409–421 (2009).

12. B. Jia, R. Serra-Moreno, W. Neidermyer, A. Rahmberg, J. Mackey, I. B. Fofana, W. E. Johnson, S. Westmoreland, D. T. Evans, Species-specific activity of SIV Nef and HIV-1 Vpu in overcoming restriction by tetherin/BST2. PLoS pathogens 5, e1000429 (2009).

13. C. Prelli Bozzo, A. Laliberté, A. De Luna, C. Pastorio, K. Regensburger, S. Krebs, A. Graf, H. Blum, M. Volcic, K. M. J. Sparrer, F. Kirchhoff, Replication competent HIV-guided CRISPR screen identifies antiviral factors including targets of the accessory protein Nef. Nat Commun 15, 3813 (2024).

14. K. Sato, N. Misawa, J. S. Takeuchi, T. Kobayashi, T. Izumi, H. Aso, S. Nagaoka, K. Yamamoto, I. Kimura, Y. Konno, Y. Nakano, Y. Koyanagi, Experimental Adaptive Evolution of Simian Immunodeficiency Virus SIVcpz to Pandemic Human Immunodeficiency Virus Type 1 by Using a Humanized Mouse Model. Journal of Virology 92, 10.1128/jvi.01905-17 (2018).

15. F. Bibollet-Ruche, A. Heigele, B. F. Keele, J. L. Easlick, J. M. Decker, J. Takehisa, G. Learn, P. M. Sharp, B. H. Hahn, F. Kirchhoff, Efficient SIVcpz replication in human lymphoid tissue requires viral matrix protein adaptation. Journal of Clinical Investigation 122, 1644–1652 (2012).

16. R. T. Abraham, A. Weiss, Jurkat T cells and development of the T-cell receptor signalling paradigm. Nat Rev Immunol 4, 301–308 (2004).

17. R. E. Means, T. Matthews, J. A. Hoxie, M. H. Malim, T. Kodama, R. C. Desrosiers, Ability of the V3 Loop of Simian Immunodeficiency Virus To Serve as a Target for Antibody-Mediated Neutralization: Correlation of Neutralization Sensitivity, Growth in Macrophages, and Decreased Dependence on CD4. Journal of Virology 75, 3903–3915 (2001).

18. P. Lusso, F. Cocchi, C. Balotta, P. D. Markham, A. Louie, P. Farci, R. Pal, R. C. Gallo, M. S. Reitz, Growth of macrophage-tropic and primary human immunodeficiency virus type 1 (HIV-1) isolates in a unique CD4+ T-cell clone (PM1): failure to downregulate CD4 and to interfere with cell-line-tropic HIV-1. Journal of Virology 69, 3712–3720 (1995).

19. M. Hsu, J. M. Harouse, A. Gettie, C. Buckner, J. Blanchard, C. Cheng-Mayer, Increased mucosal transmission but not enhanced pathogenicity of the CCR5-tropic, simian AIDS-inducing simian/human immunodeficiency virus SHIV(SF162P3) maps to envelope gp120. J Virol 77, 989–998 (2003).

20. X. Wei, J. M. Decker, H. Liu, Z. Zhang, R. B. Arani, J. M. Kilby, M. S. Saag, X. Wu, G. M. Shaw, J. C. Kappes, Emergence of resistant human immunodeficiency virus type 1 in patients receiving fusion inhibitor (T-20) monotherapy. Antimicrobial agents and chemotherapy 46, 1896–905 (2002).

21. P. Sagulenko, V. Puller, R. A. Neher, TreeTime: Maximum-likelihood phylodynamic analysis. Virus Evol 4, vex042 (2018).

22. F. Kirchhoff, H. W. Kestler, R. C. Desrosiers, Upstream U3 sequences in simian immunodeficiency virus are selectively deleted in vivo in the absence of an intact nef gene. J Virol 68, 2031–2037 (1994).

23. J. Münch, D. Rajan, E. Rücker, S. Wildum, N. Adam, F. Kirchhoff, The role of upstream U3 sequences in HIV-1 replication and CD4+ T cell depletion in human lymphoid tissue ex vivo. Virology 341, 313– 320 (2005).

24. F. Kirchhoff, T. C. Greenough, D. B. Brettler, J. L. Sullivan, R. C. Desrosiers, Absence of Intact nef Sequences in a Long-Term Survivor with Nonprogressive HIV-1 Infection. New England Journal of Medicine 332, 228–232 (1995).

25. P. J. McLaren, A. Gawanbacht, N. Pyndiah, C. Krapp, D. Hotter, S. F. Kluge, N. Götz, J. Heilmann, K. Mack, D. Sauter, D. Thompson, J. Perreaud, A. Rausell, M. Munoz, A. Ciuffi, F. Kirchhoff, A. Telenti, Identification of potential HIV restriction factors by combining evolutionary genomic signatures with functional analyses. Retrovirology 12, 41 (2015).

26. D. Mahé, G. Matusali, C. Deleage, R. L. L. S. Alvarenga, A.-P. Satie, A. Pagliuzza, R. Mathieu, S. Lavoué, B. Jégou, L. R. de França, N. Chomont, L. Houzet, A. D. Rolland, N. Dejucq-Rainsford, Potential for Virus Endogenization in Humans through Testicular Germ Cell Infection: the Case of HIV. J Virol 94, e01145–20 (2020).

27. Z. Yuan, G. Kang, L. Daharsh, W. Fan, Q. Li, SIVcpz closely related to the ancestral HIV-1 is less or non-pathogenic to humans in a hu-BLT mouse model. Emerg Microbes Infect 7, 59 (2018).

28. D. M. Tebit, G. Nickel, R. Gibson, M. Rodriguez, N. J. Hathaway, K. Bain, A. L. Reyes-Rodriguez, P. Ondoa, J. L. Heeney, Y. Li, J. Bongorno, D. Canaday, D. McDonald, J. A. Bailey, E. J. Arts, Replicative fitness and pathogenicity of primate lentiviruses in lymphoid tissue, primary human and chimpanzee cells: relation to possible jumps to humans. eBioMedicine 100, 104965 (2024).

29. W. Li, H. Xu, T. Xiao, L. Cong, M. I. Love, F. Zhang, R. A. Irizarry, J. S. Liu, M. Brown, X. S. Liu, MAGeCK enables robust identification of essential genes from genome-scale CRISPR/Cas9 knockout screens. Genome Biol 15, 554 (2014).

30. A. Rattani, M. Wolna, M. Ploquin, W. Helmhart, S. Morrone, B. Mayer, J. Godwin, W. Xu, O. Stemmann, A. Pendas, K. Nasmyth, Sgol2 provides a regulatory platform that coordinates essential cell cycle processes during meiosis I in oocytes. eLife 2, e01133 (2013).

31. K. Mardinian, J. J. Adashek, G. P. Botta, S. Kato, R. Kurzrock, SMARCA4: Implications of an altered chromatin-remodeling gene for cancer development and therapy. Mol Cancer Ther, 10.1158/1535-7163.MCT-21-0433 (2021).

32. D. L. Burdette, K. M. Monroe, K. Sotelo-Troha, J. S. Iwig, B. Eckert, M. Hyodo, Y. Hayakawa, R. E. Vance, STING is a direct innate immune sensor of cyclic di-GMP. Nature 478, 515–518 (2011).

33. S. Ikeda, S. Kishida, H. Yamamoto, H. Murai, S. Koyama, A. Kikuchi, Axin, a negative regulator of the Wnt signaling pathway, forms a complex with GSK-3beta and beta-catenin and promotes GSK-3beta-dependent phosphorylation of beta-catenin. EMBO J 17, 1371–1384 (1998).

34. J. Dutrieux, D. M. Portilho, N. J. Arhel, U. Hazan, S. Nisole, TRIM5α is a SUMO substrate. Retrovirology 12, 28 (2015).

35. N. Selliah, M. Zhang, S. White, P. Zoltick, B. E. Sawaya, T. H. Finkel, R. Q. Cron, FOXP3 inhibits HIV-1 infection of CD4 T-cells via inhibition of LTR transcriptional activity. Virology 381, 161–167 (2008).

36. D. Holmes, G. Knudsen, S. Mackey-Cushman, L. Su, FoxP3 Enhances HIV-1 Gene Expression by Modulating NFκB Occupancy at the Long Terminal Repeat in Human T Cells. J Biol Chem 282, 15973– 15980 (2007).

37. S. Iordanskiy, Y. Zhao, L. Dubrovsky, T. Iordanskaya, M. Chen, D. Liang, M. Bukrinsky, Heat Shock Protein 70 Protects Cells from Cell Cycle Arrest and Apoptosis Induced by Human Immunodeficiency Virus Type 1 Viral Protein R. J Virol 78, 9697–9704 (2004).

38. P. Chaudhary, S. Z. Khan, P. Rawat, T. Augustine, D. A. Raynes, V. Guerriero, D. Mitra, HSP70 binding protein 1 (HspBP1) suppresses HIV-1 replication by inhibiting NF-κB mediated activation of viral gene expression. Nucleic Acids Res 44, 1613–1629 (2016).

39. R. Sugiyama, H. Nishitsuji, A. Furukawa, M. Katahira, Y. Habu, H. Takeuchi, A. Ryo, H. Takaku, Heat shock protein 70 inhibits HIV-1 Vif-mediated ubiquitination and degradation of APOBEC3G. J Biol Chem 286, 10051–10057 (2011).

40. M. Lusic, B. Marini, H. Ali, B. Lucic, R. Luzzati, M. Giacca, Proximity to PML Nuclear Bodies Regulates HIV-1 Latency in CD4+ T Cells. Cell Host and Microbe 13, 665–677 (2013).

41. J. Hokello, A. Lakhikumar Sharma, M. Tyagi, AP-1 and NF-κB synergize to transcriptionally activate latent HIV upon T-cell receptor activation. FEBS Lett 595, 577–594 (2021).

42. A. G. Lloyd, S. Tateishi, P. D. Bieniasz, M. A. Muesing, M. Yamaizumi, L. C. F. Mulder, Effect of DNA repair protein Rad18 on viral infection. PLoS Pathog 2, e40 (2006).

43. C. E. Jones, W. S. Tan, F. Grey, D. J. Y. 2021 Hughes, Discovering antiviral restriction factors and pathways using genetic screens. Journal of General Virology 102, 001603.

44. S. Brezgin, A. Kostyusheva, E. Bayurova, E. Volchkova, V. Gegechkori, I. Gordeychuk, D. Glebe, D. Kostyushev, V. Chulanov, Immunity and Viral Infections: Modulating Antiviral Response via CRISPR– Cas Systems. Viruses 13, 1373 (2021).

45. S. Amini-Bavil-Olyaee, Y. J. Choi, J. H. Lee, M. Shi, I. C. Huang, M. Farzan, J. U. Jung, The antiviral effector IFITM3 disrupts intracellular cholesterol homeostasis to block viral entry. Cell Host and Microbe 13, 452–464 (2013).

46. K. Li, R. M. Markosyan, Y. M. Zheng, O. Golfetto, B. Bungart, M. Li, S. Ding, Y. He, C. Liang, J. C. Lee, E. Gratton, F. S. Cohen, S. L. Liu, IFITM Proteins Restrict Viral Membrane Hemifusion. PLoS Pathogens 9 (2013).

47. T. L. Foster, H. Wilson, S. S. Iyer, K. Coss, K. Doores, S. Smith, P. Kellam, A. Finzi, P. Borrow, B. H. Hahn, S. J. D. Neil, Resistance of Transmitted Founder HIV-1 to IFITM-Mediated Restriction. Cell Host and Microbe 20, 429–442 (2016).

48. A. A. Compton, T. Bruel, F. Porrot, A. Mallet, M. Sachse, M. Euvrard, C. Liang, N. Casartelli, O. Schwartz, IFITM proteins incorporated into HIV-1 virions impair viral fusion and spread. Cell Host and Microbe 16, 736–747 (2014).

49. J. Liu, Q. Xiao, J. Xiao, C. Niu, Y. Li, X. Zhang, Z. Zhou, G. Shu, G. Yin, Wnt/β-catenin signalling: function, biological mechanisms, and therapeutic opportunities. Sig Transduct Target Ther 7, 1–23 (2022).

50. S. Klein, G. Golani, F. Lolicato, C. Lahr, D. Beyer, A. Herrmann, M. Wachsmuth-Melm, N. Reddmann, R. Brecht, M. Hosseinzadeh, A. Kolovou, J. Makroczyova, S. Peterl, M. Schorb, Y. Schwab, B. Brügger, W. Nickel, U. S. Schwarz, P. Chlanda, IFITM3 blocks influenza virus entry by sorting lipids and stabilizing hemifusion. Cell Host & Microbe 31, 616–633.e20 (2023).

51. K. Tartour, R. Appourchaux, J. Gaillard, X.-N. Nguyen, S. Durand, J. Turpin, E. Beaumont, E. Roch, G. Berger, R. Mahieux, D. Brand, P. Roingeard, A. Cimarelli, IFITM proteins are incorporated onto HIV-1 virion particles and negatively imprint their infectivity. Retrovirology 11, 103 (2014).

52. J. Yu, S.-L. Liu, The Inhibition of HIV-1 Entry Imposed by Interferon Inducible Transmembrane Proteins Is Independent of Co-Receptor Usage. Viruses 10, 413 (2018).

53. M. Agarwal, K. K. Lai, I. Wilt, S. Majdoul, A. A. Jolley, M. Lewinski, A. A. Compton, Restriction of HIV-1 infectivity by interferon and IFITM3 is counteracted by Nef. Sci Adv 11, eadz7083 (2025).

54. L. J. Henderson, L. Al-Harthi, Role of β-Catenin/TCF-4 Signaling in HIV Replication and Pathogenesis: Insights to Informing Novel Anti-HIV Molecular Therapeutics. J Neuroimmune Pharmacol 6, 10.1007/s11481-011-9266–7 (2011).

55. H. J. Barbian, M. S. Seaton, S. D. Narasipura, J. Wallace, R. Rajan, B. E. Sha, L. Al-Harthi, β-catenin regulates HIV latency and modulates HIV reactivation. PLoS Pathog 18, e1010354 (2022).

56. D.-L. Dai, C. Xie, L.-Y. Zhong, S.-X. Liu, L.-L. Zhang, H. Zhang, X.-P. Wu, Z.-M. Wu, K. Kang, Y. Li, Y.-M. Sun, T.-L. Xia, C.-S. Zhang, A. Zhang, M. Shi, C. Sun, M.-L. Chen, G.-X. Zhao, G.-L. Bu, Y.-T. Liu, K.-Y. Huang, Z. Zhao, S.-X. Li, X.-Y. Zhang, Y.-F. Yuan, S.-J. Wen, L. Zhang, B.-K. Li, Q. Zhong, M.-S. Zeng, AXIN1 boosts antiviral response through IRF3 stabilization and induced phase separation. Signal Transduct Target Ther 9, 281 (2024).

57. J. Yang, H. Xu, F. Shao, Immunological function of familial Mediterranean fever disease protein Pyrin. Sci China Life Sci 57, 1156–1161 (2014).

58. P. Broz, V. M. Dixit, Inflammasomes: mechanism of assembly, regulation and signalling. Nat Rev Immunol 16, 407–420 (2016).

59. F. Wouters, J. Bogie, A. Wullaert, J. van der Hilst, Recent Insights in Pyrin Inflammasome Activation: Identifying Potential Novel Therapeutic Approaches in Pyrin-Associated Autoinflammatory Syndromes. J Clin Immunol 44, 8 (2023).

60. K. M. J. Sparrer, S. Gableske, M. A. Zurenski, Z. M. Parker, F. Full, G. J. Baumgart, J. Kato, G. Pacheco-Rodriguez, C. Liang, O. Pornillos, J. Moss, M. Vaughan, M. U. Gack, TRIM23 mediates virus-induced autophagy via activation of TBK1. Nature microbiology 2, 1543–1557 (2017).

61. S. Lukasiak, A. Kalinka, N. Gupta, A. Papadopoulos, K. Saeed, U. McDermott, G. J. Hannon, D. Ross-Thriepland, D. Walter, A benchmark comparison of CRISPRn guide-RNA design algorithms and generation of small single and dual-targeting libraries to boost screening efficiency. BMC Genomics 26, 198 (2025).

62. X. Yao, Z. Liu, X. Wang, Y. Wang, Y.-H. Nie, L. Lai, R. Sun, L. Shi, Q. Sun, H. Yang, Generation of knock-in cynomolgus monkey via CRISPR/Cas9 editing. Cell Res 28, 379–382 (2018).

63. M. Yolamanova, C. Meier, A. K. Shaytan, V. Vas, C. W. Bertoncini, F. Arnold, O. Zirafi, S. M. Usmani, J. A. Müller, D. Sauter, C. Goffinet, D. Palesch, P. Walther, N. R. Roan, H. Geiger, O. Lunov, T. Simmet, J. Bohne, H. Schrezenmeier, K. Schwarz, L. Ständker, W.-G. Forssmann, X. Salvatella, P. G. Khalatur, A. R. Khokhlov, T. P. J. Knowles, T. Weil, F. Kirchhoff, J. Münch, Peptide nanofibrils boost retroviral gene transfer and provide a rapid means for concentrating viruses. Nature Nanotechnology 8, 130–136 (2013).

64. L. Koepke, B. Winter, A. Grenzner, K. Regensburger, S. Engelhart, J. A. van der Merwe, S. Krebs, H. Blum, F. Kirchhoff, K. M. J. Sparrer, An improved method for high-throughput quantification of autophagy in mammalian cells. Sci Rep 10, 12241 (2020).

